# Traumatic brain injury-induced fear generalization in mice involves hippocampal memory trace dysfunction and is alleviated by (*R,S*)-ketamine

**DOI:** 10.1101/2023.02.24.529876

**Authors:** Josephine C. McGowan, Liliana R. Ladner, Claire X. Shubeck, Juliana Tapia, Christina T. LaGamma, Amanda Anqueira-González, Ariana DeFrancesco, Briana K. Chen, Holly C. Hunsberger, Ezra J. Sydnor, Ryan W. Logan, Tzong-Shiue Yu, Steven G. Kernie, Christine A. Denny

## Abstract

**INTRODUCTION:** Traumatic brain injury (TBI) is a debilitating neurological disorder caused by an impact to the head by an outside force. TBI results in persistent cognitive impairments, including fear generalization, the inability to distinguish between aversive and neutral stimuli. The mechanisms underlying fear generalization have not been fully elucidated, and there are no targeted therapeutics to alleviate this symptom of TBI.

**METHODS:** To identify the neural ensembles mediating fear generalization, we utilized the ArcCreER^T2^ x enhanced yellow fluorescent protein (EYFP) mice, which allow for activity-dependent labeling and quantification of memory traces. Mice were administered a sham surgery or the controlled cortical impact (CCI) model of TBI. Mice were then administered a contextual fear discrimination (CFD) paradigm and memory traces were quantified in numerous brain regions. In a separate group of mice, we tested if (*R,S*)-ketamine could decrease fear generalization and alter the corresponding memory traces in TBI mice.

**RESULTS:** TBI mice exhibited increased fear generalization when compared with sham mice. This behavioral phenotype was paralleled by altered memory traces in the DG, CA3, and amygdala, but not by alterations in inflammation or sleep. In TBI mice, (*R,S*)-ketamine facilitated fear discrimination and this behavioral improvement was reflected in DG memory trace activity.

**CONCLUSIONS:** These data show that TBI induces fear generalization by altering fear memory traces, and that this deficit can be improved with a single injection of (*R,S*)-ketamine. This work enhances our understanding of the neural basis of TBI-induced fear generalization and reveals potential therapeutic avenues for alleviating this symptom.

## INTRODUCTION

Traumatic brain injury (TBI) is described as the “silent epidemic” as many cases go unrecognized due to their complex, often subjective, diagnoses of the injury [1]. Despite high rates of underdiagnosis, TBI accounts for most of the death and disability worldwide among all trauma-related injuries [2,3]. TBI is a heterogeneous disorder with a complex, multi-stage pathology, yielding primary and secondary injury sequela [4]. TBI is most often the consequence of rapid acceleration or deceleration, a penetrating object, blast waves, or blunt force, where each unique instance initiates a diverse cascade of neurobiological pathways and neuropsychological symptoms. Individuals with TBI frequently experience mechanobiological shifts in the brain milliseconds after the insult, whereas deficits in attention, executive function, memory, and behavior could last for months post-injury [5].

Post-traumatic stress disorder (PTSD) is often comorbid with TBI, especially in the military where 62% of veterans who sustain multiple head injuries also experience PTSD [6]. A principal cognitive symptom observed after the onset of TBI/PTSD is fear generalization, or the maladaptive generalization of fear from an adverse context to a neutral context [7]. The inability to distinguish threatening and safe environments leads to maladaptive fear expression in contexts not directly related to the past trauma. However, the neural mechanisms underlying TBI-induced fear generalization are not well understood, and there exist no targeted therapeutics to treat this symptom. In mice, fear generalization can be quantified using a contextual fear discrimination (CFD) paradigm, which tests fear behavior in aversive and similar, but neutral contexts [7]. CFD is partially mediated by the dorsal portion of the hippocampus (HPC) [8]. The whole HPC supports learning and the storage of remote episodic memories while the dorsal HPC is specifically implicated in cognitive functions including spatial learning [8,9]. Contextual fear learning also requires coordinated activity between the HPC and the amygdala (AMG) [10,11]. Projections from the HPC to the basal AMG (BA) and central AMG (CeA) play a key role in associative learning of contextual fear [10,12,13].

To identify individual fear memory traces, numerous laboratories have utilized immediate early genes (IEGs), which are rapidly activated in response to cellular stimuli or even upon learning [14]. Our laboratory previously engineered the ArcCreER^T2^ x enhanced yellow fluorescent protein (EYFP) mouse line to permit indelible labeling of activity-regulated cytoskeleton associate protein (*Arc*/*Arg3*.*1)*^+^ cells. The permanently labeled EYFP^+^ cells can then be compared with recently activated Arc^+^ or c-fos^+^ cells to quantify individual memory traces throughout the brain [15]. We have previously shown that fear memories are encoded and stored in the dentate gyrus (DG) and Cornu Ammonis 3 (CA3) subregions of the HPC with the ArcCreER^T2^ x EYFP mice [16].

Here, we report that using a controlled cortical impact (CCI) model of TBI, TBI resulted in an increased fear generalization phenotype – specifically, sham mice could discriminate between an aversive and neutral context, but TBI mice could not discriminate between these two similar contexts. This behavioral phenotype was paralleled by altered memory traces in the DG, CA3, and AMG. While sham mice exhibited a significant difference in memory trace reactivation after exposures to the aversive and neutral contexts, TBI mice exhibited similar levels of memory trace reactivation in the DG, paralleling the CFD overgeneralization phenotype. In addition to memory traces, here, we probed whether TBI-induced fear generalization may also be related to disturbances in sleep, a common symptom in patients with TBI [17-19], or inflammation in the hippocampus. Here, sleep-wake disturbances only occurred during light-dark transitions in TBI mice, while there was no change in hippocampal inflammation. These data indicated that sleep and inflammation are not robust

To our knowledge, no study to date has identified novel therapeutics to alleviate the fear generalization phenotype seen in TBI. Previous studies from our lab have reported that a single dose of (*R,S*)-ketamine administered prior to stress is effective as a prophylactic against stress-induced behavioral despair [20], in buffering learned fear [21,22], and in decreasing fear generalization [23]. Given (*R,S*)-ketamine’s robust resilience-enhancing effects in various models of stress, we hypothesized that (*R,S*)-ketamine may also be beneficial against fear generalization in a mouse model of TBI. In TBI mice, we also report that a single dose of (*R,S*)-ketamine (30 mg/kg) administered 1 hour after a CCI facilitated fear discrimination and this behavioral improvement was reflected in DG memory trace activity. Overall, these data suggest that TBI predisposes the brain to dysregulated activation of fear-associated HPC and amygdalar memory traces and that administering (*R,S*)-ketamine facilitates fear discrimination in TBI mice by targeting DG memory traces. This work enhances our understanding of the neural basis of TBI-induced fear generalization and reveals potential therapeutic avenues for alleviating this symptom.

## METHODS AND MATERIALS

### Mice

129S6/SvEv male mice were purchased from Taconic (Hudson, NY) at 7-8 weeks of age. ArcCreER^T2^ [16] x R26R-STOP-floxed-EYFP [24] homozygous female mice were bred with R26R-STOP-floxed EYFP homozygous male mice in-house. All experimental mice were ArcCreER^T2^(+) and homozygous for the EYFP reporter and are on a 129S6/SvEv background. All mice were housed 4-5 per cage in a 12 h (06:00-18:00 h) light-dark colony room at 22°C. Food and water were provided *ad libitum*. Behavioral testing was conducted during the light phase. All mice utilized for behavioral experiments were approximately 9-12 weeks of age at the start of the experiment. All procedures were approved by the Institutional Animal Care and Use Committee (IACUC) at the New York State Psychiatric Institute (NYSPI).

### Controlled Cortical Impact (CCI) Surgery

A moderate CCI surgery was performed with a standard protocol, as previously described with a slight modification [25]. Because the current study used 129S6/SvEv and not C57BL/6J mice as previously utilized, the injury was generated with a 3 mm stainless steel tipped impactor, set to an injury depth of 0.9 mm (instead of 0.7 mm) and impact velocity of 4.4 m/s to similarly mimic a moderate TBI. Sham-treated animals were similarly anesthetized and incised, without craniectomy or cortical injury. Mice were randomly assigned to surgery groups.

**All other Methods are listed in the Supplement**.

## RESULTS

### TBI increases behavioral despair, but does not alter avoidance behavior

We first sought to characterize the effect of a moderate CCI [26] on mood and behavior in the ArcCreER^T2^ x EYFP mice. TBI was confirmed by hippocampal deformation and cortical atrophy (**Figure S1**). Mice were administered a sham or CCI surgery and 1 month later, behavioral assays were administered to quantify behavioral despair and avoidance behavior (**Figure 1A**). On day 1 of the forced swim test (FST), sham and TBI mice exhibited comparable immobility time (**Figure S2A-S2B**). On day 2 of the FST, TBI mice had significantly increased average immobility time when compared to sham mice (**Figure S2C-S2D**). In the elevated plus maze (EPM), both groups spent a comparable amount of time in the open (**Figure S2E-S2F**) and closed (**Figure S2G-S2H**) arms. These data indicate that CCI increases behavioral despair without affecting avoidance behavior.

**Figure 1.**
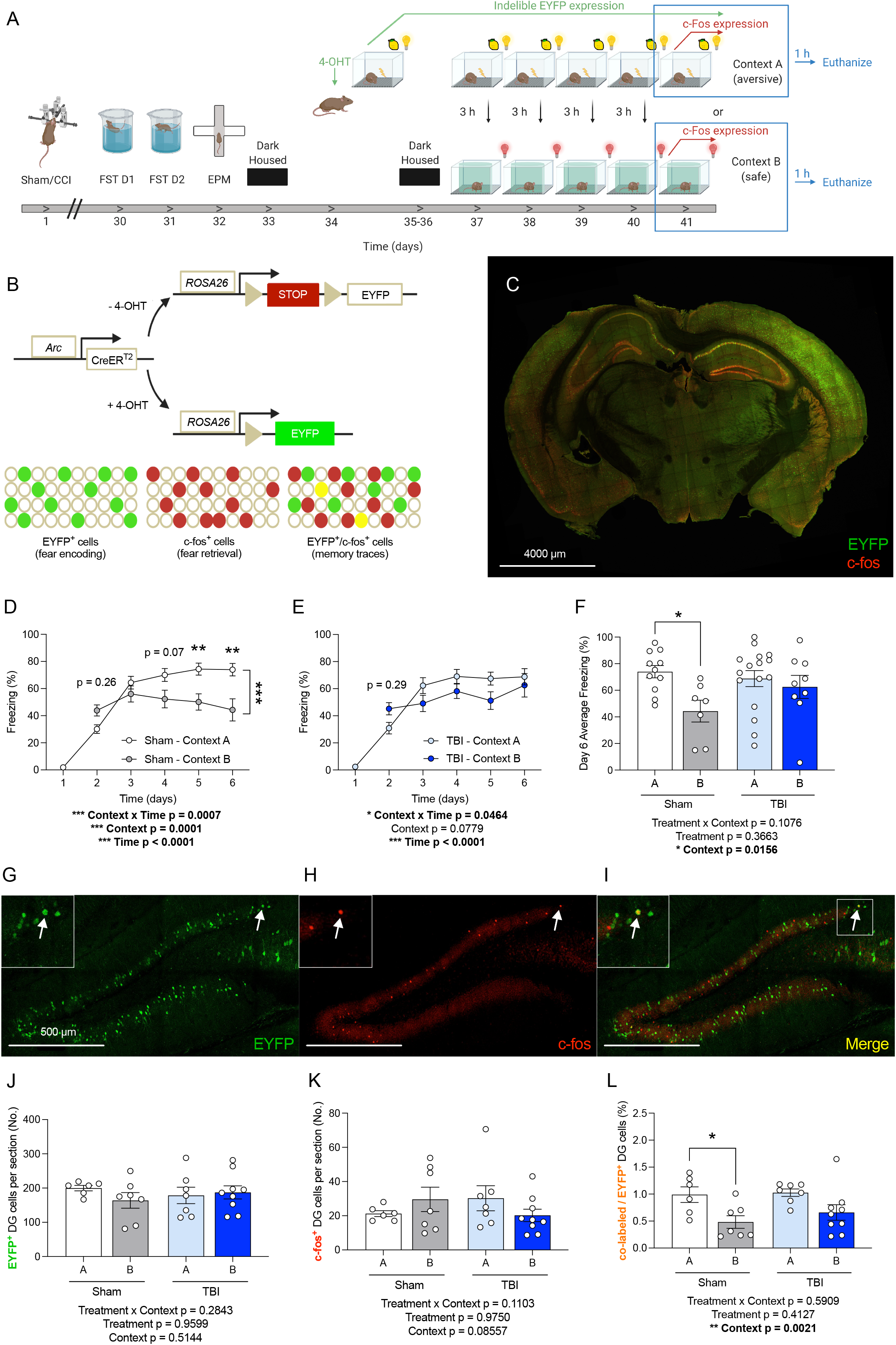
TBI impairs CFD and alters DG memory trace activity. (**A**) Experimental design. (**B**) Overview of ArcCreER^T2^ x EYFP transgenic mouse model. (**C**) Brain demonstrating sufficient cortical impact. All TBI brains exhibited cortical atrophy and hippocampal deformation, indicative of a moderate CCI. (**D**) On Days 4-6 of the CFD task, sham mice exhibited less freezing in the neutral Context B when compared to the aversive Context A. (**E**) TBI mice exhibited similar freezing in the aversive A and neutral B contexts. (**G**) On Day 6 of the CFD task, sham mice exhibited less freezing in the neutral Context B as compared to the aversive Context A whereas TBI mice exhibited similar average freezing in both contexts. (**G**) EYFP^+^ cells in the DG. (**H**) c-fos^+^ cells in the DG. (**I**) Overlap of EYFP^+^ and c-fos^+^ cells in the DG. Arrows indicate yellow co-labeled cells. (**J**) EYFP^+^ DG cells active during encoding did not differ across groups. (**K**) c-fos^+^ cells active during retrieval did not differ across groups. (**L**) Sham mice had less percentage of co-labeled/EYFP^+^ DG cells in Context B when compared with Context A. Conversely, TBI mice had comparable percentage of co-labeled/EYFP^+^ DG cells in both contexts. (n = 9-16 male mice per group). Error bars represent ± SEM. * p < 0.05, ** p < 0.01, and *** p < 0.001. TBI, traumatic brain injury; CCI, controlled cortical impact; CFD, contextual fear discrimination; EYFP, enhanced yellow fluorescent protein; μm, micrometers; No., number; DG, dentate gyrus.

### TBI increases fear generalization and alters the corresponding DG memory traces

Next, in order to tag the neural ensembles mediating fear generalization, ArcCreER^T2^ x EYFP mice were injected with 4-OHT 5 hours before the start of a CFD (**Figure 1A-1C**). Five days after the initial 1-shock contextual fear conditioning (CFC) training, mice were exposed daily to Context A (i.e., aversive context) and Context B (i.e., neutral, but similar context). Whereas sham mice were able to discriminate between the two contexts by the 6^th^ day (**Figure 1D**), TBI mice could not discriminate (**Figure 1E**). Sham mice froze significantly less in context B than in context A, but this difference was absent in TBI mice (**Figure 1F**). These data suggest that the CCI model of TBI results in increased fear generalization in the ArcCreER^T2^ x EYFP mice.

Since the dorsal DG has been implicated in CFD [7,23,27], we next quantified dorsal DG memory traces in ArcCreER^T2^ x EYFP mice (**Figures 1G-1I**). On the 6^th^ day of CFD, mice were euthanized 1 h after the last exposure to either Context A or Context B. The number of EYFP^+^ (**Figures 1J**) and c-fos^+^ (**Figures 1K**) DG cells did not differ between the groups. However, there was a significant effect of Context on the percentage of co-labeled/EYFP^+^ DG cells (**Figure 1L**). Sham mice exhibited significantly less co-labeled/EYFP^+^ DG cells in Context B than in context A. However, TBI mice exhibited comparable co-labeled/EYFP^+^ DG cells in both contexts. These data suggest that TBI alters DG memory traces, and this effect parallels the fear overgeneralization phenotype seen in TBI mice.

### TBI alters neural activity in CA3, but not in a context-dependent manner

Our prior memory trace work in the ArcCreER^T2^ x EYFP mice has shown that memory traces of contextual fear memories are encoded and stored not only in the DG, but also CA3 [16].

Moreover, previous computational and experimental evidence demonstrate that CA3 plays a role in pattern completion, whereas the DG is critical for pattern [28-30]. Therefore, we next quantified memory traces in CA3 (**Figure 2A-2C**). There was a significant effect of Treatment on the number of EYFP^+^ (**Figure 2D**) and c-fos^+^ (**Figure 2E**) CA3 cells. TBI mice exhibited fewer of these cells when compared to sham mice. There was also a significant effect of Group on the percentage of co-labeled/EYFP^+^ CA3 cells (**Figure 2F**). TBI mice exhibited a greater percentage of co-labeled/EYFP^+^ CA3 cells when compared to sham mice. These data suggest that the increased fear generalization phenotype may be driven by the increased memory trace reactivation in CA3, resulting in aberrant pattern completion in TBI mice.

**Figure 2.**
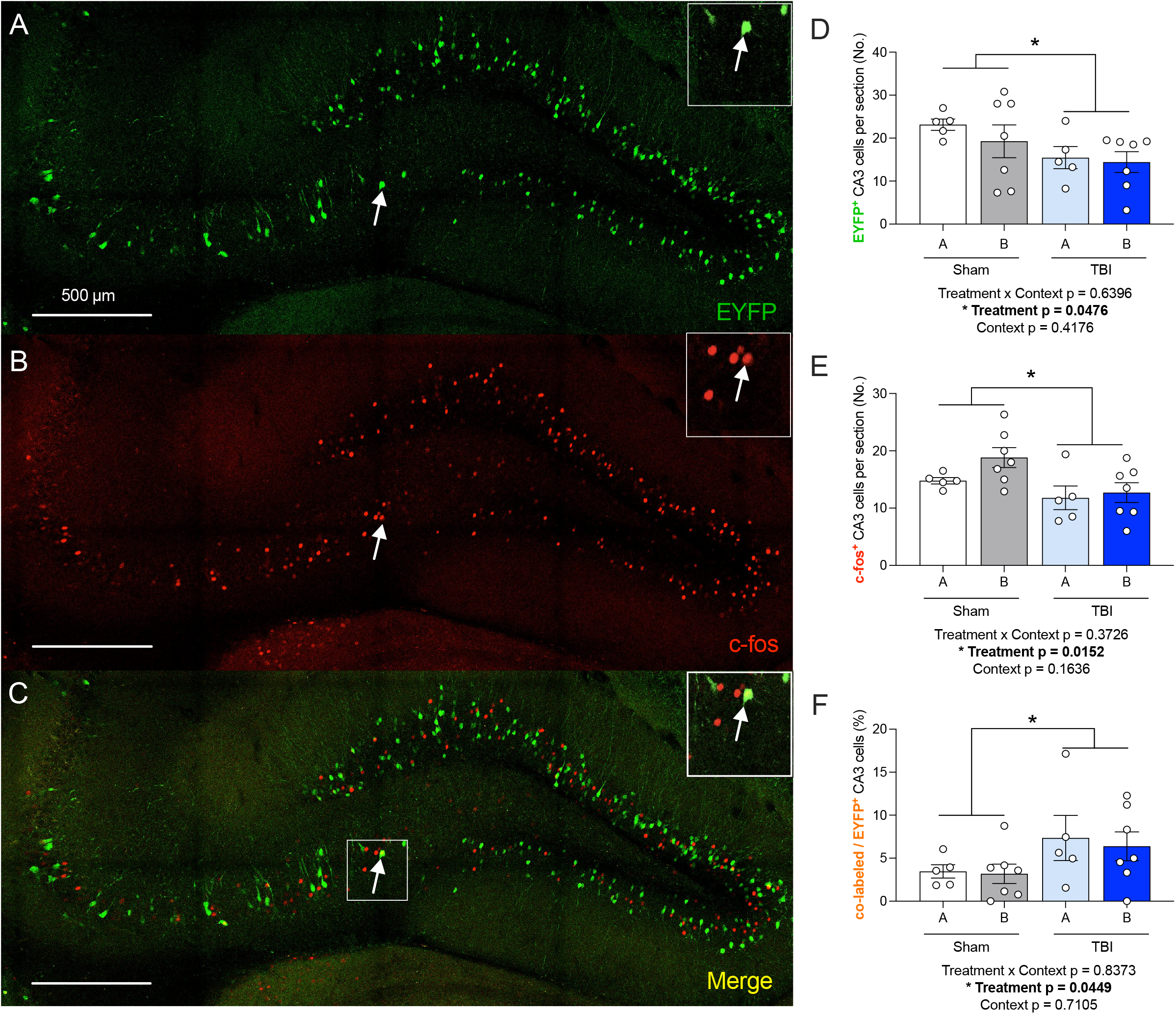
TBI increases CA3 memory trace activity. (**A**) EYFP^+^ cells in CA3. (**B**) c-fos^+^ cells in CA3. (**C**) Overlap of EYFP^+^ and c-fos^+^ cells in CA3. Arrows indicate yellow co-labeled cells. (**D**) There were fewer EYFP^+^ CA3 cells in TBI mice as compared to sham mice. (**E**) There were significantly fewer c-fos^+^ CA3 cells in TBI mice as compared to sham mice. (**F**) However, TBI mice had higher percentages of co-labeled/EYFP^+^ CA3 cells in both Contexts A and B as compared to Sham mice. (n = 5-7 male mice per group). Error bars represent ± SEM. * p < 0.05. EYFP, enhanced yellow fluorescent protein; μm, micrometers; CA3, *cornu ammonis* 3; No., number.

### TBI alters neural activity in amygdala subregions

Finally, we quantified memory traces in two subregions of the AMG, the BA and CeA, as these are the two primary subregions of the AMG that are essential for expression of fear (**Figures 3A-C**) [31-32]. In the BA, there were no significant differences in the number of EYFP^+^ (**Figure 3D**) or c-fos^+^ (**Figure 3E**) cells between the groups. However, there was a significant effect of Context, Treatment, and a Context x Treatment interaction on the percentage of co-labeled/EYFP^+^ BA cells (**Figure 3F**). In sham mice, there were significantly less co-labeled/EYFP^+^ cells in mice exposed to context B when compared with context A. This effect was not observed in TBI mice; moreover, TBI mice exhibited a significantly decreased percentage of co-labeled/EYFP^+^ BA cells.

**Figure 3.**
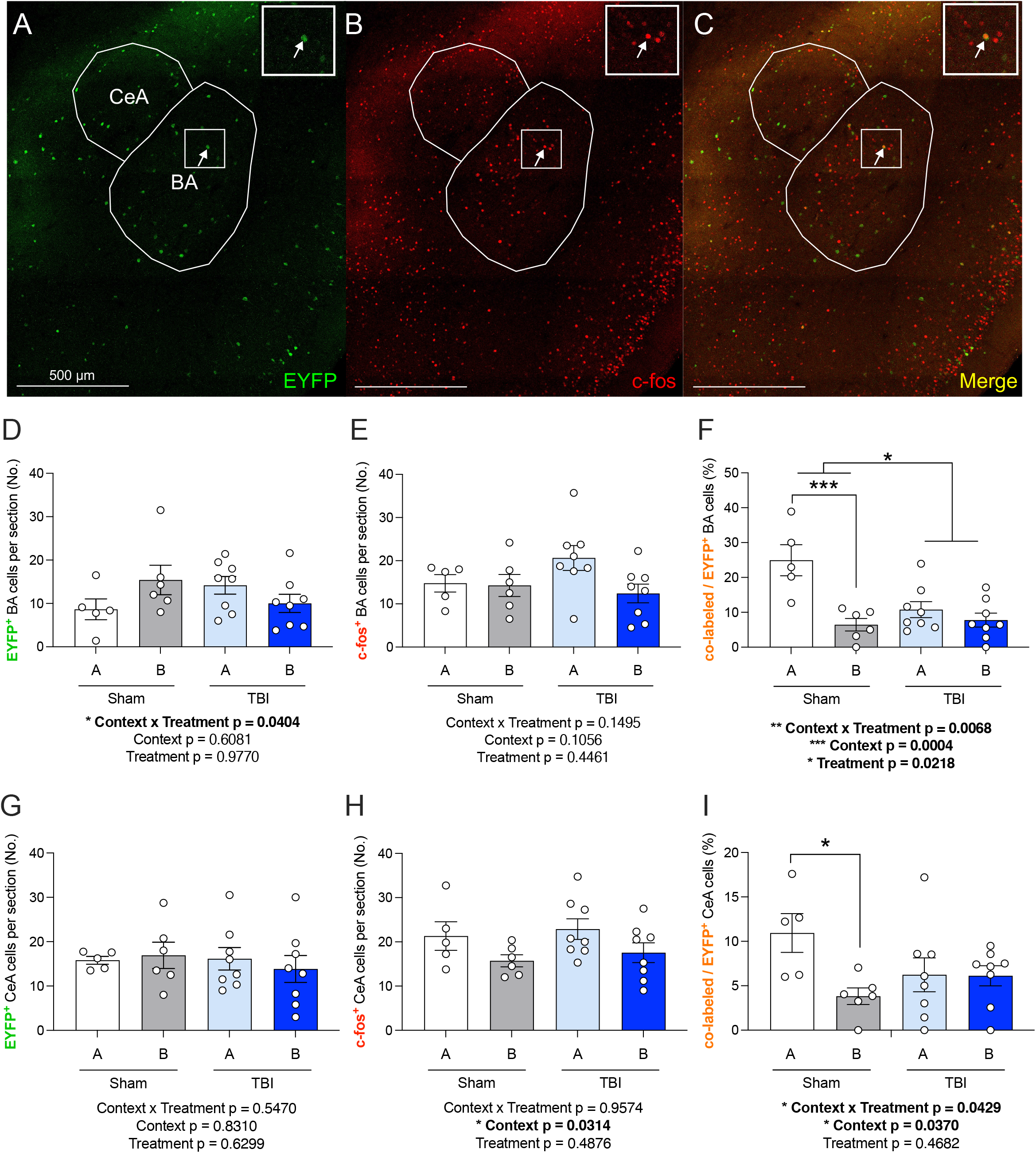
TBI alters amygdalar memory trace activity. (**A**) EYFP^+^ cells in the BA and CeA. (**B**) c-fos^+^ cells in the BA and CeA. (**C**) Example cells showing overlap of an EYFP^+^ and c-fos^+^ cell. Arrows indicate yellow co-labeled cells. (**D**) EYFP^+^ BA cells active during encoding did not differ across groups. (**E**) c-fos^+^ BA cells active during retrieval did not differ across groups. (**F**) There are significantly fewer co-labeled/EYFP^+^ BA cells in sham mice sacrificed after context B as compared to sham mice sacrificed after context A. Conversely, TBI mice showed no difference in the percentage of co-labeled/EYFP^+^ BA cells between contexts A and B. Additionally, TBI mice expressed significantly fewer co-labeled/EYFP^+^ BA cells overall as compared to sham mice. (**G**) EYFP^+^ CeA cells active during encoding did not differ across groups. (**H**) c-fos^+^ CeA cells active during retrieval did not differ across groups. (**I**) There are significantly fewer co-labeled/EYFP^+^ CeA cells in sham mice sacrificed after context B as compared to sham mice sacrificed after context A. Conversely, TBI mice showed no difference in the percentage of co-labeled/EYFP^+^ CeA cells between contexts A and B. (n = 5-8 male mice per group). Error bars represent ± SEM. * p < 0.05, ** p < 0.01, and *** p < 0.001. EYFP, enhanced yellow fluorescent protein; TBI, traumatic brain injury; CCI, controlled cortical impact; CFD, contextual fear discrimination; BA, basal amygdala; CeA, central amygdala; μm, micrometers; No., number.

In the CeA, there were no significant differences in the number of EYFP^+^ (**Figure 3G**) or c-fos^+^ (**Figure 3H**) cells between the groups. However, there was a significant effect of Context and a significant interaction of Context x Treatment on the percentage of co-labeled/EYFP^+^ CeA cells (**Figure 3I**). In sham mice, there were significantly less co-labeled/EYFP^+^ cells in mice exposed to context B when compared with context A. This effect was not observed in TBI mice. In summary, these data suggest that in addition to alterations in HPC memory traces, TBI mice induce significant alterations in AMG memory traces that parallel behavioral deficits.

### TBI does not result in chronic inflammatory changes in the hippocampus

To determine whether factors beyond memory trace reactivation may be contributing to TBI-induced fear generalization, hippocampal inflammation was assessed, as neuroinflammation is one of the hallmark symptoms of TBI [33] and is the major cause of secondary cell death following TBI [34]. Brains obtained for the memory trace experiment (**Figure S3A**) were additionally processed for glial fibrillary acidic protein (GFAP) (**Figure S3B**) and cyclooxygenase-2 (Cox-2) (**Figure S3C-S3D**). Surprisingly, there were no differences in expression of GFAP (**Figure S4E-S4F**) or Cox-2 (**Figure S3G-S4H**) in either the DG or CA3. These data indicate that inflammatory processes in the HPC likely do not significantly contribute to the long-term behavioral deficits in TBI mice.

### TBI minimally impacts sleep architecture

Since TBI is a risk factor for development of sleep and circadian rhythm disruptions [17-19], we next sought to determine whether changes in sleep architecture may also contribute to TBI-induced behavioral alterations. We measured sleep-wake changes before and after a 3-shock CFC paradigm using non-invasive sleep monitoring systems. A sham or CCI procedure was first performed on the mice (**Figure 4A**). Sleep was then assessed at 1-week post-impact. Mice were then administered CFC at the 1-month timepoint, and sleep was assessed for the 2^nd^ time starting at 42 days post-impact. During CFC training, there was no effect of TBI on fear encoding (data not shown). However, during context re-exposure, TBI mice exhibited increased freezing when compared to sham mice (**Figure 4B**).

**Figure 4.**
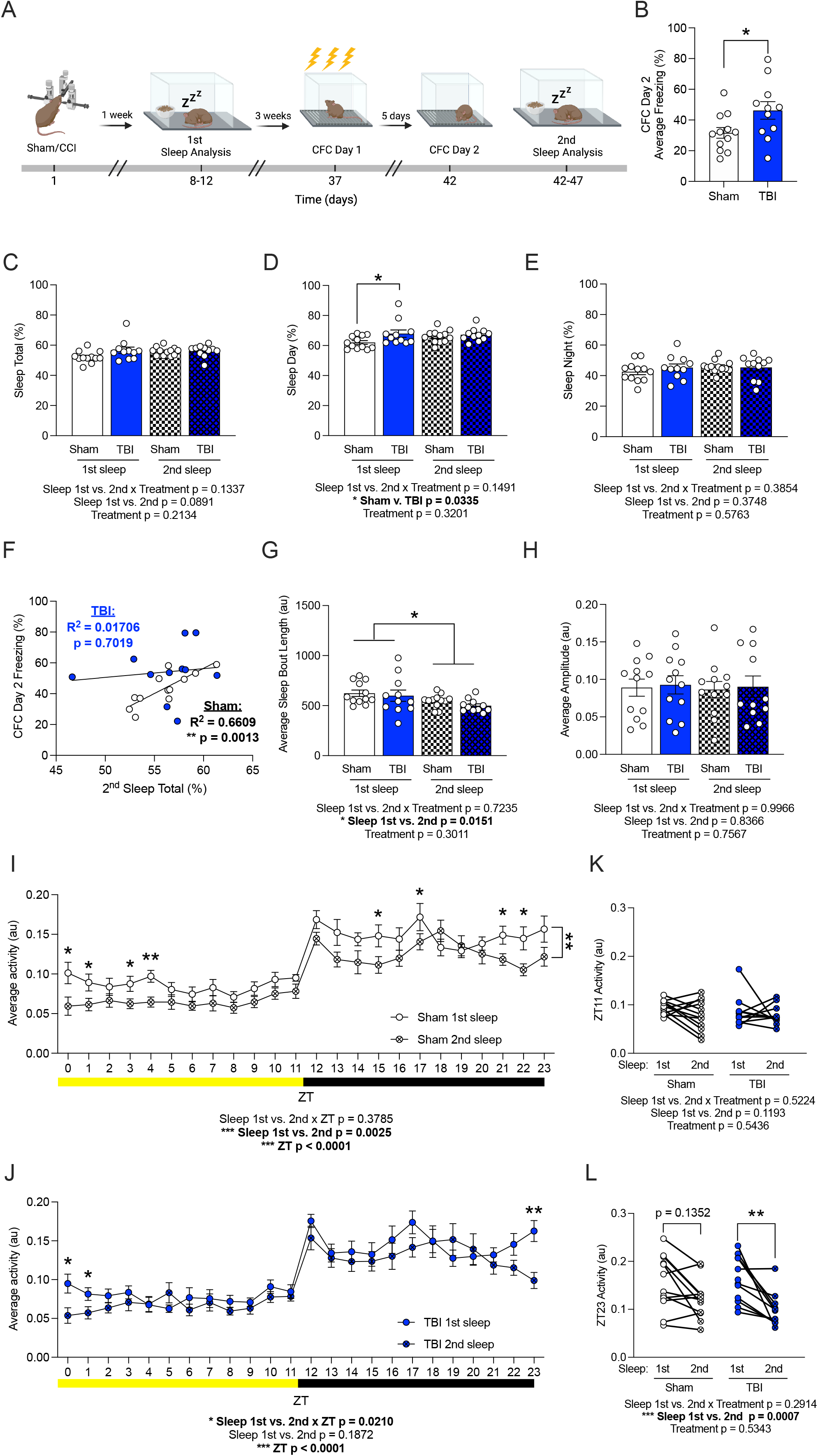
TBI increases fear behavior and minimally alters sleep architecture. (**A**) Experimental design. (**B**) Average freezing during day 2 of the CFC was significantly higher in TBI mice as compared to sham mice. (**C**) There was no difference in total sleep between sham or TBI mice across the 1^st^ and 2^nd^ sleep. (**D**) Only during the 1^st^ sleep, TBI mice slept more during the day than sham mice. (**E**) There was no difference in sleep during the night between sham or TBI mice across the 1^st^ and 2^nd^ sleep. (**F**) In sham but not TBI mice, total sleep during the 2^nd^ sleep assessment positively correlated with freezing on day 2 of the CFC. (**G**) Average sleep bout length decreased in both sham and TBI mice from the 1^st^ to the 2^nd^ sleep. (**H**) Average sleep amplitude did not differ across groups. (**I**) Sham mice exhibit significantly lower average activity across 24 h during the 2^nd^ sleep as compared to the 1^st^ sleep. (**J**) TBI mice exhibited significantly lower average activity only during ZT23-1 from the 1^st^ to the 2^nd^ sleep. (**K**) There was no change in activity from the 1^st^ to the 2^nd^ sleep during ZT11 across groups. (**L**) Activity during ZT23 was significantly decreased from the 1^st^ to the 2^nd^ sleep only in TBI mice. (n = 11-12 male mice per group). Error bars represent ± SEM. * p < 0.05, ** p < 0.01. TBI, traumatic brain injury; EPM, elevated plus maze; FST, forced swim test; NSF; novelty suppressed feeding; CFC, contextual fear conditioning; sec, seconds; min, minute; au, arbitrary unites; ZT, Zeitgeber time.

All groups exhibited a comparable percentage of total sleep (**Figure 4C**). However, there was a significant effect of Treatment on the percentage of day sleep (**Figure 4D**). During the 1^st^ sleep analysis, TBI mice displayed increased sleep during the day as compared to sham mice, an effect which was not observed during the 2^nd^ sleep (**Figure 4D**). All groups exhibited a comparable percentage of night sleep (**Figure 4E**).

We next sought to determine whether freezing during day 2 of CFC correlated with the percentage of sleep during the 2^nd^ sleep analysis. Interestingly, sham, but not TBI, mice displayed a positive correlation between total sleep (%) during the 2^nd^ sleep and freezing (%) (**Figure 4F**). To determine if there are other aspects of sleep that may be altered uniquely in TBI mice, the average sleep bout length and amplitude of sleep was next quantified. While the average sleep bout length decreased from the 1^st^ to 2^nd^ sleep, there was no effect of Treatment (**Figure 4G**). All groups of mice exhibited a comparable amplitude (**Figure 4H**).

We next assessed diurnal activity changes from the 1^st^ to the 2^nd^ sleep assessments by analyzing changes across Zeitgeber time (ZT). In sham mice, there was a significant effect of the 1^st^ vs. 2^nd^ sleep session and of ZT on average activity across time; sham mice exhibited decreased activity during the 2^nd^ sleep session when compared with the 1^st^ session (**Figure 4I, Figure S4A-S4D**). In TBI mice, there was a significant effect of ZT, but not of the 1^st^ vs. 2^nd^ sleep session (**Figure 4J, Figure S4E-S4H**). However, there was a significant interaction of 1^st^ vs. 2^nd^ sleep x ZT; average activity decreased during the 2^nd^ sleep session when compared with the 1^st^ only during ZT23-1 (i.e., during the night to morning transition) (**Figure 4J**). To determine if the transition changes were altered in TBI mice, we assessed activity specifically during ZT11 (light to dark) and ZT23 (dark to light) (**Figures 4K-4L**). While there was no change in activity from the first to second sleep in both sham and TBI mice during ZT11 (**Figure 4K**), activity during ZT23 was only decreased in TBI mice from the first to second sleep (**Figure 4L**). In both sham and TBI mice, there was comparable amplitude between the 1^st^ and 2^nd^ sessions (**Figure S4I-L**). Overall, these data suggest that although TBI mice exhibited increased fear expression, sleep architecture is only minimally affected in the CCI model.

### (*R,S*)-ketamine facilitates fear discrimination in TBI mice, which is reflected in DG memory trace activity

Finally, to determine if TBI-induced fear generalization may potentially be alleviated, a single dose of (*R,S*)-ketamine (30 mg/kg) was administered 1 h post-CCI to ArcCreER^T2^ x EYFP mice (**Figure 5A**). One h post-impact was chosen as a clinically relevant timepoint, as a single administration of a drug can feasibly be implemented as part of the regimen for patients with a recent TBI. One month later, the FST, EPM, and CFD paradigms were administered. On both days 1 and 2 of the FST, there was no difference between the groups (**Figures S5A-S5D**). Additionally, time spent in the open or closed arms of the EPM did not differ between groups (**Figures S5E-S5H**).

**Figure 5.**
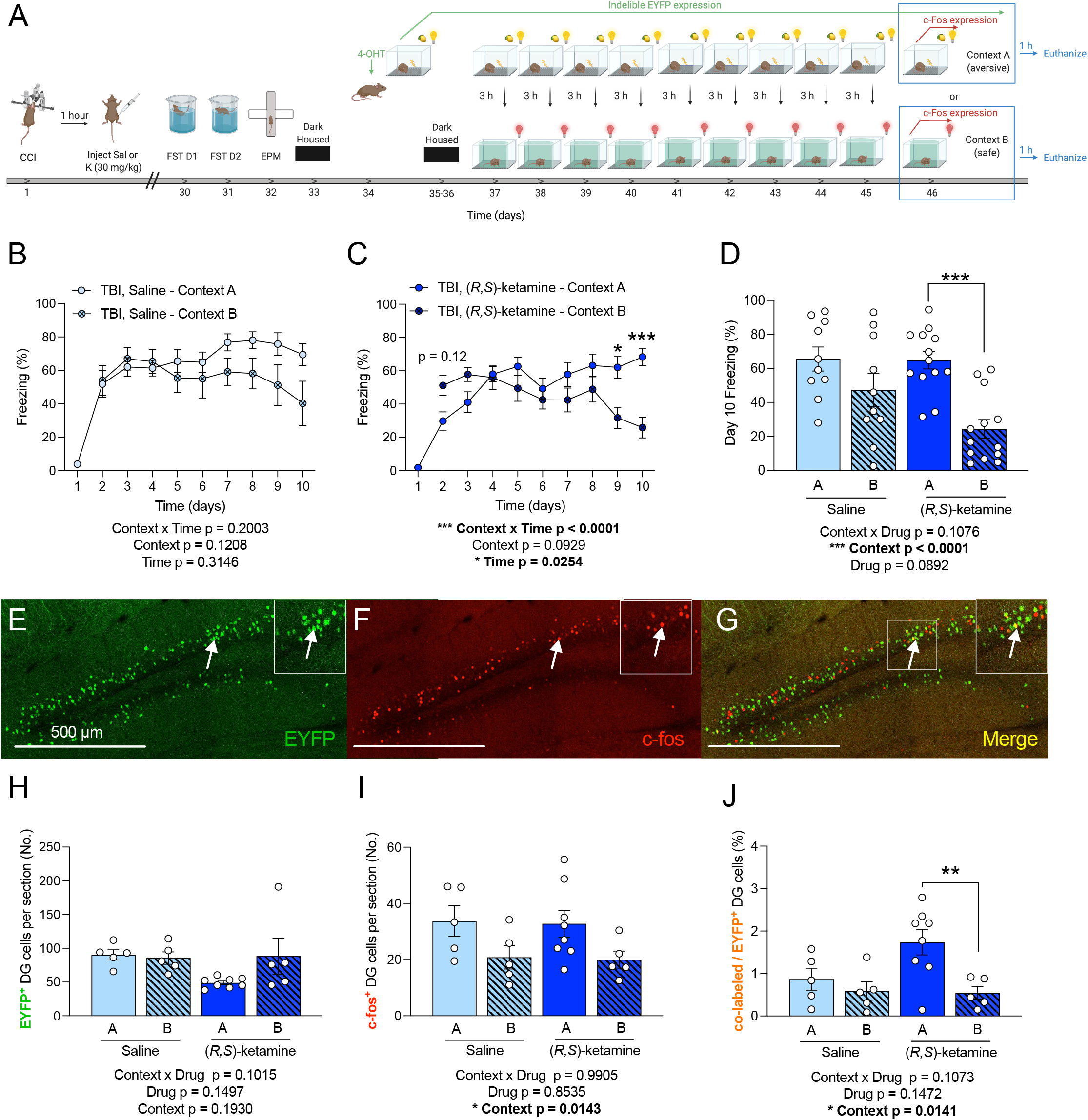
Post-impact (*R,S*)-ketamine administration facilitates fear discrimination in TBI mice which is reflected in DG memory trace activity. (**A**) Experimental design. (**B**) Saline-treated TBI mice did not differ in freezing between the neutral Context B as compared to the aversive Context A across all days. (**C**) On Days 9-10 of the CFD task, (*R,S*)-ketamine-treated TBI mice exhibited significantly more freezing in the aversive Context A as compared to the neutral Context B. (**D**) Freezing (%) on Day 10 of the CFD task did not significantly differ between Contexts A and B in saline-treated TBI mice, but (*R,S*)-ketamine-treated mice demonstrated significantly increased freezing in context A as compared to context B. (**E**) EYFP^+^ cells in the DG. (**F**) c-fos^+^ cells in the DG. (**G**) Overlap of EYFP^+^ and c-fos^+^ cells in the DG. Arrows indicate yellow co-labeled cells. (**H**) EYFP^+^ DG cells active during encoding did not differ across groups. (**I**) c-fos^+^ DG cells active during retrieval did not differ across groups. (**J**) Saline-treated TBI mice had comparable percentage of co-labeled/EYFP^+^ DG cells in contexts A and B. However, (*R,S*)-ketamine-treated TBI mice demonstrated increased percentage of co-labeled/EYFP^+^ DG cells in Context A when compared with Context B. (n = 10-13 male mice per group). Error bars represent ± SEM. * p < 0.05, ** p < 0.01, and *** p < 0.001. CCI, controlled cortical impact; TBI, traumatic brain injury; FST, forced swim test; EPM, elevated plus maze; CFD, contextual fear discrimination; 4-OHT, 4-hydroxytamoxifen; EYFP, enhanced yellow fluorescent protein; μm, micrometers; DG, dentate gyrus; No., number; h, hours; Sal, saline; K, (*R,S*)-ketamine (30 mg/kg).

In the CFD task, while saline-treated TBI mice did not discriminate (**Figure 5B**), (*R,S*)-ketamine-treated TBI mice froze significantly less in Context B as compared to Context A during days 9 and 10 of the task, demonstrating facilitated fear discrimination (**Figure 5C-5D**).

To assess whether the behavioral effects seen following (*R,S*)-ketamine administration are reflected in activated neural ensembles, we next quantified DG memory traces (**Figure 5E-5G**). The number of EYFP^+^ DG cells did not differ between the groups (**Figure 5H**). However, there was an overall effect of Context, but not Drug, on the number of c-fos^+^ DG cells (**Figure 5I**). Interestingly, while saline-treated TBI mice had comparable percentages of co-labeled/EYFP^+^ DG cells, (*R,S*)-ketamine-treated TBI exhibited a significant difference between mice exposed to context A and context B (**Figure 5J**) - the percentage of co-labeled/EYFP^+^ cells in was significantly less in context B than in Context A. These data indicate that post-impact (*R,S*)-ketamine administration facilitates CFD partially via DG memory trace activity, revealing a potential node to target to alleviate TBI-induced fear generalization.

## DISCUSSION

Here, we examined the effects of the CCI model of TBI on fear overgeneralization in the activity dependent ArcCreER^T2^ x EYFP mice. TBI increased fear generalization and this behavioral phenotype was paralleled by altered hippocampal and amygdalar fear memory traces. Finally, in TBI mice, a single dose of (*R,S*)-ketamine facilitated fear discrimination, and altered the corresponding DG memory traces. Overall, these data reveal that a core symptom of TBI, fear generalization, can be successfully modeled and pharmacologically modulated in a CCI rodent model of TBI.

The present study behaviorally assessed mice in the FST, EPM, and CFD 1 month after TBI, indicating that behavioral effects observed are exhibited long after an injury has occurred. The onset of depression after TBI may indeed be delayed, as depression occurs in 33-42% of TBI patients within the first year and 61% within the first 7 years following the injury [35]. One study reported that patients with moderate to severe TBI who developed mood disorders also had smaller hippocampal volumes as compared to patients with equivalent severe TBI who did not develop mood disturbance [36]. Overall, our data confirm that behavioral despair occurs well past the initial brain injury. Yet, while we found that there was no effect on avoidance behavior across all experiments, previous studies report inconsistent effects. While one study using CCI reported that TBI mice exhibited reduced avoidance behavior, their behavior was assessed 21 days post-injury [37]. Another study found no effects on avoidance behavior up to 24 days post-CCI [38]. These discrepancies suggest that the effects of TBI on avoidance behavior may be time-dependent and future studies are necessary.

We next showed that TBI resulted in impaired CFD. These results recapitulate previously observed alterations in fear learning following TBI by Reger et al. (2012), which assessed fear discrimination 2 days after a lateral fluid percussion injury model (LFPI) of TBI. As we assessed CFD 1 month after TBI, our data provide additional evidence of long-term deficits following TBI [39]. We then report that a DG fear memory trace deficit parallels this CFD behavioral effects in the ArcCreER^T2^ mice. These data are in alignment with our previous studies showing that increased fear expression is paralleled by increased levels of memory trace reactivation [16,40,42]. The fear overgeneralization phenotype observed in the TBI mice may be a result of increased reactivation of the initial traumatic memory trace not only in the aversive context but also in the neutral, similar context. Targeted modulation of this DG trace may prove to be a novel avenue for decreasing fear overgeneralization in TBI.

Interestingly, in CA3, there was an overall effect of Treatment on EYFP^+^, c-fos^+^, and co-labeled / EYFP^+^ cells; specifically, TBI mice exhibited significantly more co-labeled cells than sham mice. Because of the known role of the DG in pattern separation and CA3 in pattern completion [28-30], these data suggest that TBI biases the hippocampus towards a state of pattern completion over pattern separation, resulting in a behavioral state in which fear is generalized.

In both amygdalar subregions, we report memory trace differences in sham, but not TBI mice. We investigated the amygdala because numerous studies have shown that the hippocampus generates contextual representations that acquire associative fear expression with connections to the amygdala [13]. It is therefore possible that TBI disrupts hippocampal pathways that route contextual information to subregions of the AMG to calibrate fear responses, ultimately leading to activity that promotes fear behavior.

To determine whether inflammation may partially mediate changes in behavior and memory trace activity, levels of Cox-2 and GFAP in the hippocampal DG and CA3 were quantified. Previous studies have shown that CCI increases Cox-2 in the rat brain, but these studies only assessed Cox-2 up to several days post-impact [42,43]. Similarly, other studies have found increased GFAP expression post-injury both clinically in the serum and plasma [44] and pre-clinically in the hippocampus [45]. Here, however, we found that there were no changes in either Cox-2 or GFAP expression in the hippocampus between all groups. The lack of effect may be because brains were processed approximately 6 weeks post-CCI. However, these data are in accord with a prior study by Kernie and colleagues reporting no changes in GFAP at 60 days post-injury [25]. Thus, it is possible that hippocampal-wide inflammatory responses are limited to the initial weeks post-injury, and that at proximal times, astrogliosis occurs in defined subregions. Further studies will be necessary to elucidate the time course of inflammatory changes post-impact and whether they contribute to the fear generalization phenotype.

Since sleep-wake disturbances are one of the most prevalent symptoms of TBI [46], we next investigated how TBI might persistently alter sleep architecture. We report that TBI mice exhibited increased fear behavior in a 3-shock CFC task, which parallels the increased fear generalization observed in the CFD task. However, we did not observe major differences in sleep measurements between sham and TBI mice. The most significant effect was observed during the light cycle transition, specifically during the transition from dark to light (ZT23).

Despite this intriguing effect, the lack of significant effects in most measurements could be a result of the moderate CCI model, the relative short time that mice spend in the Piezo Sleep boxes, or the length of time following the CCI procedure. Future studies will be needed to better identify sleep disturbances in the context of TBI.

Finally, we found that administering (*R,S*)-ketamine 1 h post-impact facilitated fear discrimination in TBI mice and this behavioral improvement was paralleled by altered DG memory traces. This TBI finding is in line with our previously publication showing that (*R,S*)-ketamine can facilitate fear discrimination in adult mice [23]. Recent evidence has revealed that (*R,S*)-ketamine requires adult-born immature granule neurons in the hippocampus for its antidepressant effects [47]. It is therefore possible that (*R,S*)-ketamine targets DG adult-born granule cells during the hours post-injury to prevent alterations to intra-hippocampal circuitry, as well as projections to the AMG, eventually alleviating secondary sequelae and facilitating fear discrimination.

One of the major challenges in developing therapeutics for TBI is overcoming the blood-brain barrier (BBB) while delivering drugs [50]. However, (*R,S*)-ketamine is known to effectively cross the BBB because of its lipophilic nature [51]. There are few clinical studies investigating the efficacy of (*R,S*)-ketamine on fear behavior following TBI, as most studies were focused on successfully dispelling the myth that it increased intracranial pressure [52]. However, a recent study revealed that administering (*R,S*)-ketamine to US service members who sustained serious combat injuries related to TBI and were administered (*R,S*)-ketamine had lower incidence of PTSD outcomes during the first year postinjury as compared to those treated with opioids [53]. These data reveal that even clinically, administration of (*R,S*)-ketamine can affect fear-related behavior when administered shortly after the TBI. Future clinical studies will be necessary to determine if incorporating (*R,S*)-ketamine in the anesthetic regime for patients with moderate TBI is effective at mitigating long-term deleterious effects.

In conclusion, we found that TBI-induced deficits in CFD may be mediated by alterations to hippocampal and amygdalar fear memory traces. Administering (*R,S*)-ketamine post-impact is effective in alleviating fear generalization in TBI mice and corresponding DG memory trace deficits. Current work is focused on elucidating whether (*R,S*)-ketamine alters memory traces in other brain regions or prevent TBI-induced changes in sleep-wake disturbances. Overall, this work contributes to the understanding of neural drivers of CFD and how they become dysregulated in TBI. We offer a potential avenue for preventing long-term fear generalization in the clinic using post-injury prophylactic pharmaceutical compounds.

## ACKNOWLEDGEMENTS

JCM was supported by an NSF GRFP DGE 16-44869, an NIMH F31 MH122187-01, and an NINDS F99 NS124182. LRL was supported by the Barnard Summer Science Research Institute (SSRI). CXS was supported by Columbia University Summer Undergraduate Research Fellowship. JT was supported by the Barnard Summer Research Institute. CTL was supported by an NIH DP5 OD017908. AAG was supported by the Neurobiology & Behavior Research Training Grant T32 HD007430-19. BKC was supported by an F31 MH121023-01A1. HCH was supported by an NIA K99/R00 AG059953 Career Transition Award. EJS was supported by a retention package from the NYSPI. SGK was supported by an NIH/NINDS R56-NS-089523 and an R01-NS-095803 and the Paul Allen Foundation. CAD was supported by an NICHD R01 HD101402, an NIA R56 AG058661, an NIA R21 AG064774, and an NINDS R21 NS114870. We thank René Hen and members of the Denny and Kernie laboratories for insightful comments on this project and manuscript. Figures of behavioral timelines were created with BioRender.com.

## AUTHOR CONTRIBUTIONS

JCM, TSY, SGK, and CAD contributed to the conception and design of the work, the analysis and interpretation of the data for the work, drafting the work, and revising it critically for important intellectual content. JCM, LRL, CXS, JT, CTL, AAG, and AD ran immunohistochemistry, processed the images, quantified the cells, and performed the statistics. JCM, LRL, CXS, JT, CTL, AAG, AD, BKC, HCH, and EJS ran behavioral tests.

## REFERENCES

1. Faul, M., Xu, L., Wald, M. M. & Coronado, V. G. (ed National Center for Injury Prevention and Control Atlanta (GA): Centers for Disease Control and Prevention) (2010).

2. Dewan, M. C. et al. Estimating the global incidence of traumatic brain injury. J Neurosurg, 1–18, (2018).

3. Rubiano, A. M., Carney, N., Chesnut, R. & Puyana, J. C. Global neurotrauma research challenges and opportunities. Nature 527, S193–197 (2015).

4. Maas, A. I., Stocchetti, N. & Bullock, R. Moderate and severe traumatic brain injury in adults. Lancet Neurol 7, 728–741 (2008).

5. Somayaji, M. R., Przekwas, A. J. & Gupta, R. K. Combination Therapy for Multi-Target Manipulation of Secondary Brain Injury Mechanisms. Curr Neuropharmacol 16, 484–504 (2018).

6. Lindquist, L. K., Love, H. C. & Elbogen, E. B. Traumatic Brain Injury in Iraq and Afghanistan Veterans: New Results From a National Random Sample Study. J Neuropsychiatry Clin Neurosci 29, 254–259 (2017).

7. Sahay, A. et al. Increasing adult hippocampal neurogenesis is sufficient to improve pattern separation. Nature 472, 466–470 (2011).

8. Frankland, P. W., Cestari, V., Filipkowski, R. K., McDonald, R. J. & Silva, A. J. The dorsal hippocampus is essential for context discrimination but not for contextual conditioning. Behav Neurosci 112, 863–874 (1998).

9. Anand, K. S. & Dhikav, V. Hippocampus in health and disease: An overview. Ann Indian Acad Neurol 15, 239–246 (2012).

10. Kim, W. B., & Cho, J. H. Encoding of contextual fear memory in hippocampal-amygdala circuit. Nat Commun 11, 1382 (2020).

11. Asok, A., Kandel, E. R., & Rayman, J. B. The neurobiology of fear generalization. Front Behav Neurosci 12, 329 (2019).

12. Chaaya, N., Battle, A. R., & Johnson, L. R. An update on contextual fear memory mechanisms: Transition between amygdala and hippocampus. Neurosci Biobehav Rev 92, 43–54 (2018).

13. Tovote, P., Fadok, J. P., & Lüthi, A. Neuronal circuits for fear and anxiety. Nat Rev Neurosci 16, 317–331 (2015).

14. Denny C. A., Lebois E., & Ramirez S. From engrams to pathologies of the brain. Front Neural Circuits 11, 23 (2017).

15. Leal Santos, S. et al. Propranolol Decreases Fear Expression by Modulating Fear Memory Traces. Biological Psychiatry 89 (2021).

16. Denny, C. A. et al. Hippocampal memory traces are differentially modulated by experience, time, and adult neurogenesis. Neuron 83, 189–201 (2014).

17. Aoun, R., Rawal, H., Attarian, H., & Sahni, A. Impact of traumatic brain injury on sleep: an overview. Nat Sci Sleep 11, 131–140 (2019).

18. Duclos, C., et al. Sleep and wake disturbances following traumatic brain injury. Pathol Biol (Paris) 62, 252–261 (2014).

19. Sandsmark, D. K., Elliott, J. E., & Lim, M. M. Sleep-wake disturbances after traumatic brain injury: Synthesis of human and animal studies. Sleep 40, zsx044 (2017).

20. Brachman, R. A. et al. Ketamine as a prophylactic against stress-induced depressive-like behavior. Biological Psychiatry 79, 776–786 (2016).

21. McGowan, J. C. et al. Prophylactic ketamine attenuates learned fear. Neuropsychopharmacology 42, 1577–1589 (2017).

22. McGowan, J. C. et al. Prophylactic ketamine alters nucleotide and neurotransmitter metabolism in brain and plasma following stress. Neuropsychopharmacology 43, 1813–1821 (2018).

23. Mastrodonato, A. et al. Ventral CA3 Activation Mediates Prophylactic Ketamine Efficacy Against Stress-Induced Depressive-like Behavior. Biol Psychiatry 84, 846–856 (2018).

24. Srinivas, S. et al. Cre reporter strains produced by targeted insertion of EYFP and ECFP into the ROSA26 locus. BMC Dev Biol 1, 4 (2001).

25. Kernie, S. G., Erwin, T. M. & Parada, L. F. Brain remodeling due to neuronal and astrocytic proliferation after controlled cortical injury in mice. J Neurosci Res 66, 317–326 (2001).

26. Siebold, L., Obenaus, A., & Goyal, R. Criteria to define mild, moderate, and severe traumatic brain injury in the mouse controlled cortical impact model. Exp Neurol 310: 48–57 (2018).

27. McHugh, T. J. et al. Dentate gyrus NMDA receptors mediate rapid pattern separation in the hippocampal network. Science 317, 94–99 (2007).

28. Leutgeb, J. K., Leutbeg, S., Moser, M. B., & Moser, E. I. Pattern separation in the dentate gyrus and CA3 of the hippocampus. Science 315, 961–966 (2007).

29. Neunuebel, J. P., & Knierim, J. J. CA3 retrieves coherent representations from degraded input: direct evidence for CA3 pattern completion and dentate gyrus pattern separation. Neuron 81, 416–427.

30. Treves, A., & Rolls, E. T. Computational analysis of the role of the hippocampus in memory. Hippocampus 4, 374–391 (1994).

31. Calhoon, G.G., & Tye, K. M. Resolving the neural circuits of anxiety. Nat Neurosci 18, 1394–1404 (2015).

32. McCorkle, T. A., Barson, J. R., & Raghupathi, R. A role for the amygdala in impairments of affective behaviors following mild traumatic brain injury. 15, 601275 (2021).

33. Rovegno, M., Soto, P. A. Sáe, J. C., & von Bernhardi R. Biological mechanisms involved in the spread of traumatic brain damage. Med Intensiva 36, 37–44 (2012).

34. Schimmel, S. J., Acosta, S., & Lozano D. Neuroinflammation in traumatic brain injury: A chronic response to an acute injury. Brain Circ 3, 135–142 (2017).

35. Fann, J. R., Hart, T. & Schomer, K. G. Treatment for depression after traumatic brain injury: a systematic review. J Neurotrauma 26, 2383–2402 1091 (2009).

36. Jorge, R. E., Acion, L., Starkstein, S. E., & Magnotta, V. Hippocampal volume and mood disorders after traumatic brain injury. Biol Psychiatry 62, 332–338 (2007).

37. Washington, P. M. et al. The effect of injury severity on behavior: a phenotypic study of cognitive and emotional deficits after mild, moderate, and severe controlled cortical impact injury in mice. J Neurotrauma 29, 2283–2296 (2012).

38. Sierra-Mercado D., et al. Controlled cortical impact before or after fear conditioning does not affect fear extinction in mice. Brain Res 1606: 133–144 (2015).

39. Reger, M. L. et al. Concussive brain injury enhances fear learning and excitatory processes in the amygdala. Biol Psychiatry 71, 335–343 (2012).

40. Lacagnina, A. F. et al. Distinct hippocampal engrams control extinction and relapse of fear memory. Nat Neurosci 22, 753–761 (2019).

41. Perusini, J. N. Optogenetic stimulation of dentate gyrus engrams restores memory in Alzheimer’s disease mice. Hippocampus 27, 1110–1122 (2017).

42. Gopez, J. J., et al. Cyclooxygenase-2-specific inhibitor improves functional outcomes, provides neuroprotection, and reduces inflammation in a rat model of traumatic brain injury. Neurosurgery 56, 590–604 (2006).

43. Hickey, R. W., et al. Cyclooxygenase-2 activity following traumatic brain injury in the developing rat. Pediatr Res 62, 271–276 (2007).

44. Huebschmann, N. A., et al. Comparing glial fibrillary acidic protein (GFAP) in serum and plasma following mild traumatic brain injury in older adults. Front Neurol 11, 1054 (2020).

45. Yu, T. S. et al. Traumatic brain injury-induced hippocampal neurogenesis requires activation of early nestin-expressing progenitors. J Neurosci 28, 12901–12912 (2008).

46. Tapp, Z. M., Godbout, J. P., & Kokiko-Cochran, O. N. A tilted axis: Maladaptive inflammation and HPA axis dysfunction contribute to consequences of TBI. Front Neurol, 10: 345 (2019).

47. Rawat, R., Tunc-Ozcan, E., McGuire, T. L., Peng, C. Y., & Kessler, J. A. Ketamine activates adult-born immature granule neurons to rapidly alleviate depression-like behaviors in mice. Nat Commun 13, 2650 (2022).

48. Kanthasamy, A. et al. Chronic traumatic encephalopathy. Springer International Publishing Switzerland, Neuroimmune Pharmacology (2017).

49. Jonkman, K., Dahan, A., van de Donk, T., Aarts, L., Niesters, M., & van Velzen, M. Ketamine for pain. F1000Res 6, 1711 (2017).

50. Gregers, M. C. T., Mikkelsen, S., Lindvig, K. P., & Brochner, A. C. Ketamine as an anesthetic for patients with acute brain injury: A systematic review. Neurocrit Care 33, 273–282 (2020).

51. Melcer, T. et al. Is prehospital ketamine associated with a change in the prognosis of PTSD? Mil Med, Online ahead of print.

