## Supplemental for "Traumatic brain injury-induced fear generalization in mice involves hippocampal memory trace dysfunction and is alleviated by (*R,S*)-ketamine"

**SUPPLEMENTAL METHODS**

**Genotyping**

Genotyping was performed as previously described (Denny et al., 2014).

**Drugs**

***(R,S)-ketamine***

A single injection of saline (0.9% NaCl) or (*R,S*)-ketamine (30 mg/kg) (Ketaset III, Ketamine HCl injection, Fort Dodge Animal Health, Fort Dodge, IA) was administered once to mice at approximately 8 weeks of age. (*R,S*)-ketamine was prepared in physiological saline and all injections were administered intraperitoneally (i.p.) in volumes of 0.1 cc per 10 mg body weight.

***4-hydroxytamoxifen (4-OHT)***

Recombination was induced on Day 1 of the CFD task using 4-OHT (#H7904, Sigma-Aldrich, St. Louis, MO) as previously described (Cazzulino et al., 2015). Four-OHT was dissolved by sonication in 10% EtOH / 90% corn oil (10 mg/mL). Mice were administered 2 mg of 4-OHT via intraperitoneal (i.p.) injection.

**Behavioral Assays**

***Forced Swim Test (FST)***

The FST was administered as previously described (Brachman et al., 2016; Richardson-Jones et al., 2010). Videos were scored for mouse immobility, climbing, and swimming using an automated ViewPoint Videotrack software package (ViewPoint, Lyon, France).

***Elevated Plus Maze (EPM)***

The EPM was administered as previously described (Mallaret et al., 1999). Videos were scored by AnyMaze behavioral tracking software to measure the amount of time and number of entries into the closed arms, open arms, and center of the EPM (Stoelting, Wood Dale, IL). Time in the open arms is calculated as the sum of time in the open and center arms together.

***Contextual Fear Conditioning (CFC)***

One- and 3-shock CFC paradigms were administered as previously described (Denny et al., 2014). Briefly, mice were brought into the behavior room in a home cage, with normal lights on, and were placed in a CFC box scented with lemon (context A), to be administered 1 shock at 180s or 3 shocks at 180, 240, and 300 s after placement into the context. All sessions were scored for freezing using FreezeFrame4 (Actimetrics, Wilmette, IL).

***Contextual Fear Discrimination (CFD)***

The CFD task was administered as previously described using Contexts A and B as detailed in **Table S1** (Mastrodonato et al., 2018). For ArcCreER^T2^ experiments with no drug, mice were then anesthetized 1 h after Context A or Context B of Day 6. For the (*R,S*)-ketamine experiments, mice were anesthetized 1 h after Context A or Context B on Day 10, as these groups of mice took longer to discriminate. FreezeView4 software was used to score all sessions for freezing (Actimetrics, Wilmette, IL). Additionally, in our paradigm, ArcCreER^T2^ mice were dark housed to reduce non-specific EYFP labeling one day prior to the CFD paradigm (Denny et al., 2014). On Day 1 of the CFD paradigm, ArcCreER^T2^ mice were injected with 4-OHT to open a window of fluorescent labeling of Arc^+^ cells. Five hours after injection, ArcCreER^T2^ mice were exposed to the aversive context for the first time. Mice were then dark housed for 2 days immediately after their first exposure to the aversive context to minimize aberrant labeling of cells (Denny et al., 2014;).

***Novelty Suppressed Feeding (NSF)***

The NSF paradigm was performed as previously described (David et al., 2007). Briefly, the testing apparatus consisted of a plastic box (50 x 50 x 20 cm). The floor was covered with approximately 2 cm of wooden bedding and the arena was brightly lit (1100-1200 lux). Mice were food restricted for 18 h. All food was removed from the home cage. At the time of testing, a single pellet of food (regular chow) was placed on a white paper platform positioned in the center of the box. Each animal was placed in a corner of the box, and a stopwatch was immediately started. The latency of the mice to begin eating was recorded, with a limit of 8 minutes. Immediately after the latency was recorded, the food pellet was immediately removed from the arena. The mice were then assessed for post-restriction weight. Male and female mice were not run at the same time to avoid potential confounds. A Kaplan-Meier survival analysis was used due to the lack of normal distribution of data. The Mantel-Cox log-rank test was used to evaluate differences between the experimental groups.

**Circadian Activity and Sleep/Wake Recordings**

Circadian activity and sleep/wake measurements were determined using a non-invasive automated piezoelectric recording of movements and breath rate (PiezoSleep, Signal Solutions, LLC, Lexington, KY). Mice were individually housed 1-week post-impact in a ventilated, light-controlled room on a standard 12 h cycle of daytime light (06:00-18:00 h) with continuous data collection. The light was ∼300 lux during the light phase and ∼0 lux during the dark phase in the room. An initial 24 h acclimation period was followed by data recording for 5 d for the first sleep analysis. Mice were placed back into their home cages for behavioral testing for approximately 1 month until their second sleep analysis. Mice were randomly assigned to cages to counterbalance cage position. Each cage rested on a polyvinylidene difluoride (PVDF) square sensor (17.8 × 17.8 cm, 110 μm thick) protected by a thin plastic tray (50.8 μm) (Donohue et al., 2008). A rubber pad between each sensor and the base prevented crosstalk between the cages. The sensors were connected to an amplifier. Pressure signals and breath rates were classified as movements related to activity and inactivity or sleep and wake. Data were extracted and analyzed by using the SleepStats2p10 software (Signal Solutions, LLC, Lexington, KY) and ClockLab (Actimetrics, Wilmette, IL). Average activity per day or across all days is presented in Zeitgeber time (ZT). Amplitude is only presented for days 2-5 of sleep analysis, which contain 24 h of data.

**Tissue Collection**

Mice were anesthetized with an (*R*,*S*)-ketamine (100 mg/kg) and xylazine (10 mg/kg) mixture and transcardially perfused with 25 mL of 0.1 M phosphate buffered saline (PBS), followed by 25 mL of 4% paraformaldehyde (PFA). The brains were excised and fixed in 4% PFA overnight, then chilled in PBS at 4°C. Brains were sectioned into 100-micron serial coronal slices using a vibratome and stored in 1X PBS with 0.1% sodium azide.

**Immunohistochemistry**

To assess memory traces, sections from ArcCreER^T2^ mice were rinsed 3 times in 1X PBS, dehydrated in 50% MeOH/PBS for 2 h at room temperature, then rinsed 3 times in 0.2% phosphate buffered saline with 0.2% Tween (PBST). Blocking was performed at room temperature for 2 h in 0.2% PBST / 10% dimethyl sulfoxide (DMSO) / 6% normal donkey serum (NDS), then sections were rinsed 3 times in 0.2% PBST and 10 μg/mL heparin salt (PTwH). Incubation was performed for 3 d at 4°C in a solution of primary antibody chicken polyclonal anti-GFP (1:2000, Abcam, Cambridge, MA) and rabbit polyclonal IgG anti-c-fos (1:1000, Santa Cruz Biotechnology Inc., Dallas, TX) in PTwH / 5% DMSO / 3% NDS. Sections were then washed 3 times in PTwH and incubated overnight at 4°C in secondary antibody solution with Cy2 conjugated donkey anti-chicken IgG (1:250, Jackson ImmunoResearch, West Grove, PA) and Alexa 647 conjugated donkey anti-rabbit IgG (1:500, Life Technologies, Carlsbad, CA). Sections were rinsed 3 times in PTwH then rinsed 3 times in PBS. Sections were mounted on slides, cover slipped with Fluoromount-G, and stored at -20°C.

For inflammation experiments, sections from ArcCreER^T2^ mice were rinsed 3 times in 1X PBST, dehydrated in 50% MeOH/PBST for 2 h at room temperature, then rinsed 3 times in 0.2% PBST. Blocking was performed at room temperature for 2 h in 0.1% PBST and 10% NDS, then sections were rinsed 3 times in 0.2% PBST. Incubation was performed for 3 days at 4°C in a solution of primary antibody mouse monoclonal IgG anti-Cox-2 (1:200, Santa Cruz Biotechnology, Santa Cruz, CA) and rabbit polyclonal IgG anti-GFAP (1:500, Agilent Dako Omnis, Santa Clara, CA) in 0.1% PBST and 10% NDS. Sections were then washed in PBS 3 times and incubated overnight at 4°C in secondary antibody solution consisting of Alexa 594 conjugated donkey anti-mouse IgG (1:500, Thermo Fisher Scientific, Waltham, MA) and Cy2 conjugated donkey anti-rabbit IgG (1:250, Jackson ImmunoResearch, West Grove, PA). Sections were rinsed in PBS 3 times. Sections were mounted on slides, cover slipped with Fluoromount G, and stored at -20°C.

**Confocal Microscopy**

Samples were imaged on a confocal scanning microscope (Leica TCS SP8, Leica Microsystems Inc., Wetzlar, Germany) at 1 f/s with a dry 20X objective (NA 0.70, working distance 0.5 mm), a pixel size of 1.08 x 1.08 μm^2^, and a z step of 3.0 μm. The LAS X Matrix software-tiling algorithm was used to stitch together images of the entire slice. EYFP^+^, c-fos^+^, and co-labeled cell counts were quantified from the z-stack images by an experimenter blind to experimental groups.

For sections stained with GFAP and Cox-2, an investigator blind to treatment manually circled the granule cell layer (GCL) of the DG or pyramidal layer (PL) of CA3 throughout the entire rostro-caudal axis of the HPC using Fiji (Fiji, Dresden, Germany). The mean intensity of GFAP or Cox-2 expression was measured bilaterally using Fiji (Mastrodonato et al., 2022a; Mastrodonato et al., 2022b).

**Statistical Analysis**

The effects of Drug, Group, or Time was analyzed using an analysis of variance (ANOVA), using repeated measures where appropriate. For repeated measures ANOVA (RMANOVA), a Greenhouse-Geisser correction was implemented. Post-hoc *t*-tests and Šídák’s corrections were performed when appropriate statistical significance and interactions were achieved to warrant post-hoc analyses. For analysis of the effect of Group or Drug only on behavior, data were analyzed using unpaired two-tailed Student's *t*-test. Pearson correlations were used to compare behavior to cell count data. All data were analyzed using GraphPad Prism (v9.4.1) software (GraphPad Software Inc., La Jolla, CA). Alpha was set to 0.05 for all analyses. Data are expressed as means ± SEM. * p < 0.05, ** p < 0.01, *** p < 0.001. Main results of statistical tests are reported in the figures and more thoroughly in **Table S2**. All reagents, software, and key resources used are listed in **Table S3.**

**SUPPLEMENTAL REFERENCES**

Brachman, R. A. et al. Ketamine as a Prophylactic Against Stress-Induced Depressive-like Behavior. Biological Psychiatry 79, 776-786 (2016).

Cazzulino, A. S., Martinez, R., Tomm, N. K. & Denny, C. A. Improved specificity of hippocampal memory trace labeling. Hippocampus 26, 752-762 (2015).

David, D. J., et al. Efficacy of MCHR1 antagonist *N*-[3-(1-{[4-(3,4-Difluorophenoxy)phenyl]methyl}(4-piperidyl))-4-methylphenyl]-2-methylpropanamide (SNAP 94847) in mouse models of anxiety and depression following acute and chronic administration is independent of hippocampal neurogenesis. J Pharmacol Exp Ther 321, 237-248 (2007).

Denny, C. A. et al. Hippocampal memory traces are differentially modulated by experience, time, and adult neurogenesis. Neuron 83, 189-201 (2014).

Malleret, G., Hen, R., Guillou, J. L., Segu, L. & Buhot, M. C. 5-HT1B receptor knock-out mice exhibit increased exploratory activity and enhanced spatial memory performance in the Morris water maze. J Neurosci 19, 6157-6168 (1999).

Mastrodonato, A. et al. Ventral CA3 activation mediates prophylactic ketamine efficacy against stress-induced depressive-like behavior. Biol Psychiatry 84, 846-856 (2018).

Mastrodonato, A. et al. Acute (*R,S*)-ketamine administration induces sex-specific behavioral effects in adolescent but not aged mice. Front Neurosci 16, 852010 (2022a).

Mastrodonato, A. et al. Prophylactic (R,S)-ketamine is effective against stress-induced behaviors in adolescent but not aged mice. Int J Neuropsychopharmacol 25, 512-523 (2022b).

Richardson-Jones, J. W. et al. 5-HT1A autoreceptor levels determine vulnerability to stress and response to antidepressants. Neuron 65, 40-52 (2010).

**SUPPLEMENTAL FIGURE LEGENDS**

**Supplemental Figure 1. The CCI is an ethological, reproducible model of TBI.**
(**A**) Experimental design. A high-air pressure piston is positioned above the right cortex to induce a traumatic injury post-craniectomy. (**B**) Representative Nissl stain of a mild TBI, characterized by minimal cortical lesion and little to no hippocampal deformation. (**C**) Representative Nissl stain of a moderate TBI, characterized by cortical loss and hippocampal deformation. (**D**) Representative Nissl stain of a severe TBI, characterized by cortical loss and some hippocampal lesion. CCI, controlled cortical impact; TBI, traumatic brain injury.

**Supplemental Figure 2. TBI increases behavioral despair but does not alter anxiety-like behavior.** (**A**) On day 1 of the FST, immobility time across minutes and (**B**) average immobility time did not differ between sham or TBI mice. (**C**) On day 2 of the FST, floating duration across all minutes did not differ between sham or TBI mice. (**D**) However, during minutes 3 to 6 of Day 2 of the FST, TBI mice exhibited increased average immobility time as compared to sham mice. Across all five minutes of the EPM and on average, sham and TBI mice spent similar time in the (**E-F**) open and (**G-H**) closed arms. (n = 14-18 male mice per group). Error bars represent ± SEM. 4-OHT, 4-hydroxytamoxifen; EYFP, enhanced yellow fluorescent protein; CCI, controlled cortical impact; TBI, traumatic brain injury; FST, forced swim test; EPM, elevated plus maze; h, hour; min, minute, sec, seconds.

**Supplemental Figure 3. TBI does not alter inflammatory response in the dentate gyrus (DG) or CA3 of ArcCreER^T2^ x EYFP mice.** (**A**) Experimental design. Representative images of (**B**) GFAP and (**C**) Cox-2 staining in the hippocampus, with examples of staining in the DG. (**D**) Overlap of GFAP and Cox-2 in the hippocampus. All groups of mice expressed similar levels of GFAP in the (**E**) DG and (**F**) CA3 of the hippocampus. All groups of mice expressed similar levels of expressed similar levels of Cox-2 in the (**G**) DG and (**H**) CA3 of the hippocampus. (n = 5-8 male mice per group). Error bars represent ± SEM. EYFP, enhanced yellow fluorescent protein; TBI, traumatic brain injury; CCI, controlled cortical impact; FST, forced swim test; EPM, elevated plus maze; GFAP, glial fibrillary acidic protein; DG, dentate gyrus; CA3, *cornu ammonis* 3; Cox-2, cyclooxygenase-2; au, arbitrary units; h, hour.

**Supplemental Figure 4. TBI differentially alters sleep architecture across days.** (**A-D**) In sham mice, between the 1^st^ and 2^nd^ sleep assessments, mice displayed higher activity during the 1^st^ sleep assessment on days (**A**) 2, (**B**) 3, and (**D**) 5. (**E-H**) In TBI mice, between the 1^st^ and 2^nd^ sleep assessments, mice displayed higher activity during the 1^st^ sleep assessment only on days (**E**) 2 and (**G**) 4. (**I**) Amplitude across days between the 1^st^ and 2^nd^ sleep assessments and (**J**) average amplitude does not differ in sham mice. (**K**) Similarly, amplitude across days between the 1^st^ and 2^nd^ sleep assessments and (**L**) average amplitude does not differ in TBI mice. (n = 11-12 male mice per group). Error bars represent ± SEM. TBI, traumatic brain injury; au, arbitrary units; no., number; ZT, Zeitgeber time.

**Supplemental Figure 5. Post-impact (*R,S*)-ketamine administration does not alter behavioral despair or avoidance behavior.** (**A**) Immobility time during day 1 of the FST across all minutes and (**B**) on average does not differ between saline- or (*R,S*)-ketamine-treated TBI mice. (**C**) Immobility time during day 2 of the FST across all minutes and (**D**) on average does not differ between the groups. (**E**) Time spent in the closed arms of the EPM across time and (**F**) in total did not differ between the groups. (**G**) Time spent in the open arms of the EPM across time and (**I**) in total did not differ between the groups. CCI, controlled cortical impact; TBI, traumatic brain injury; Sal, saline; K, (*R,S*)-ketamine (30 mg/kg); FST, forced swim test; EPM, elevated plus maze; 4-OHT, 4-hydroxytamoxifen; EYFP, enhanced yellow fluorescent protein; sec, seconds; min, minutes.

**Supplemental Table 1. Information for contextual fear discrimination contexts.**

**Supplemental Table 2. Statistical analyses.**

**Supplemental Table 3. Key resources table.**
