## Supplementary figures and images for "Traumatic brain injury-induced fear generalization in mice involves hippocampal memory trace dysfunction and is alleviated by (*R,S*)-ketamine"

### Figure S1

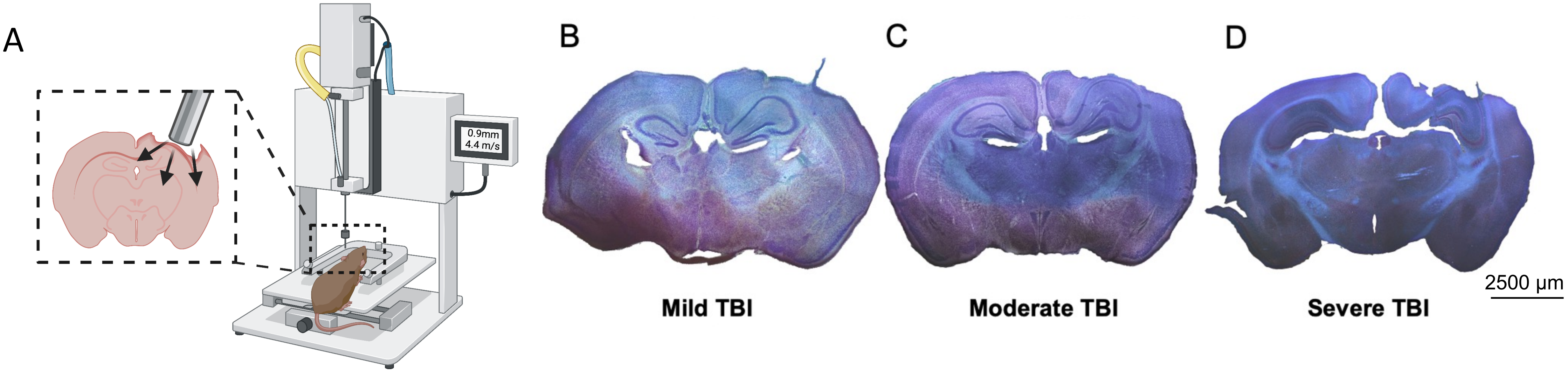

### Figure S2

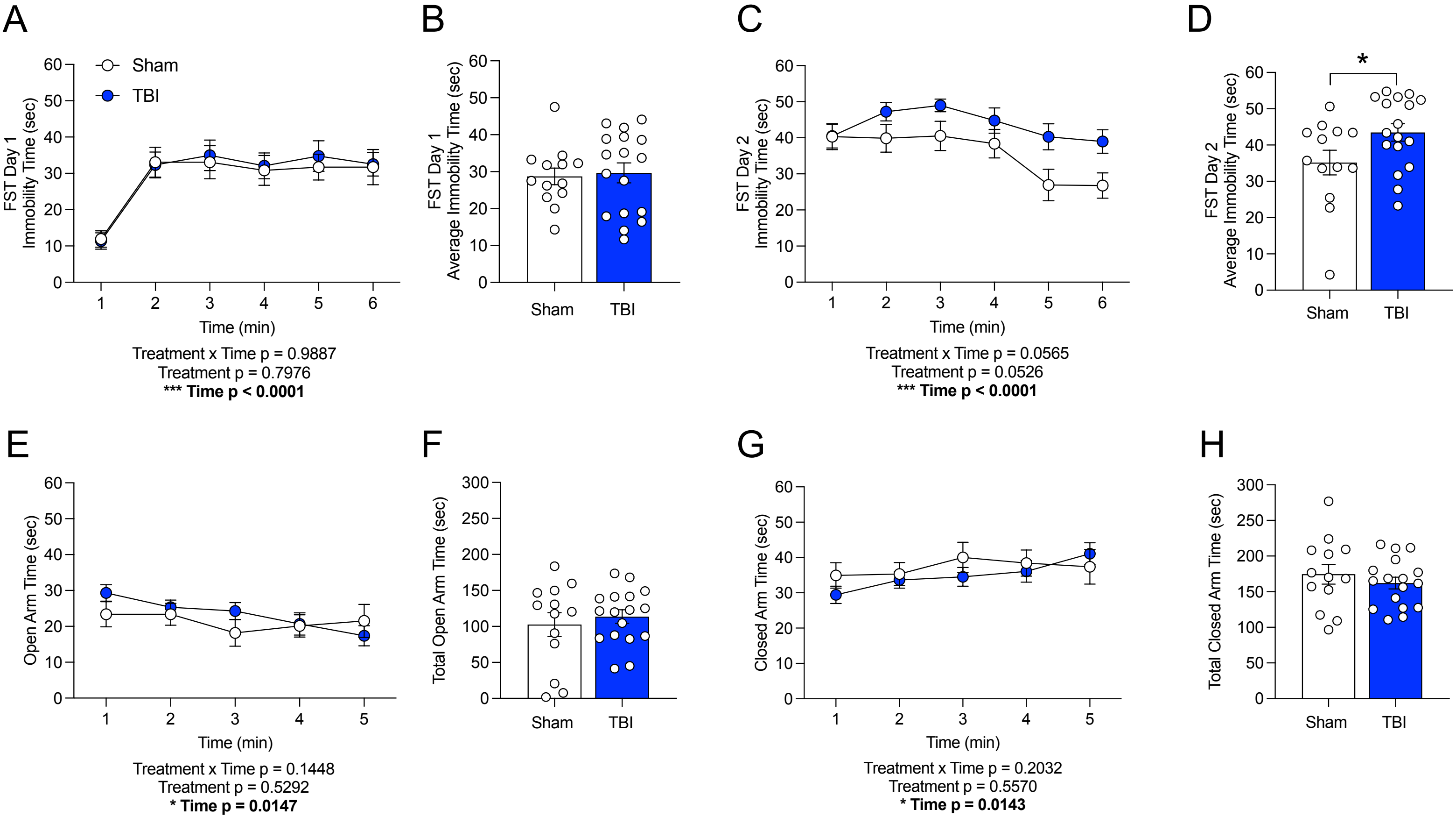

### Figure S3

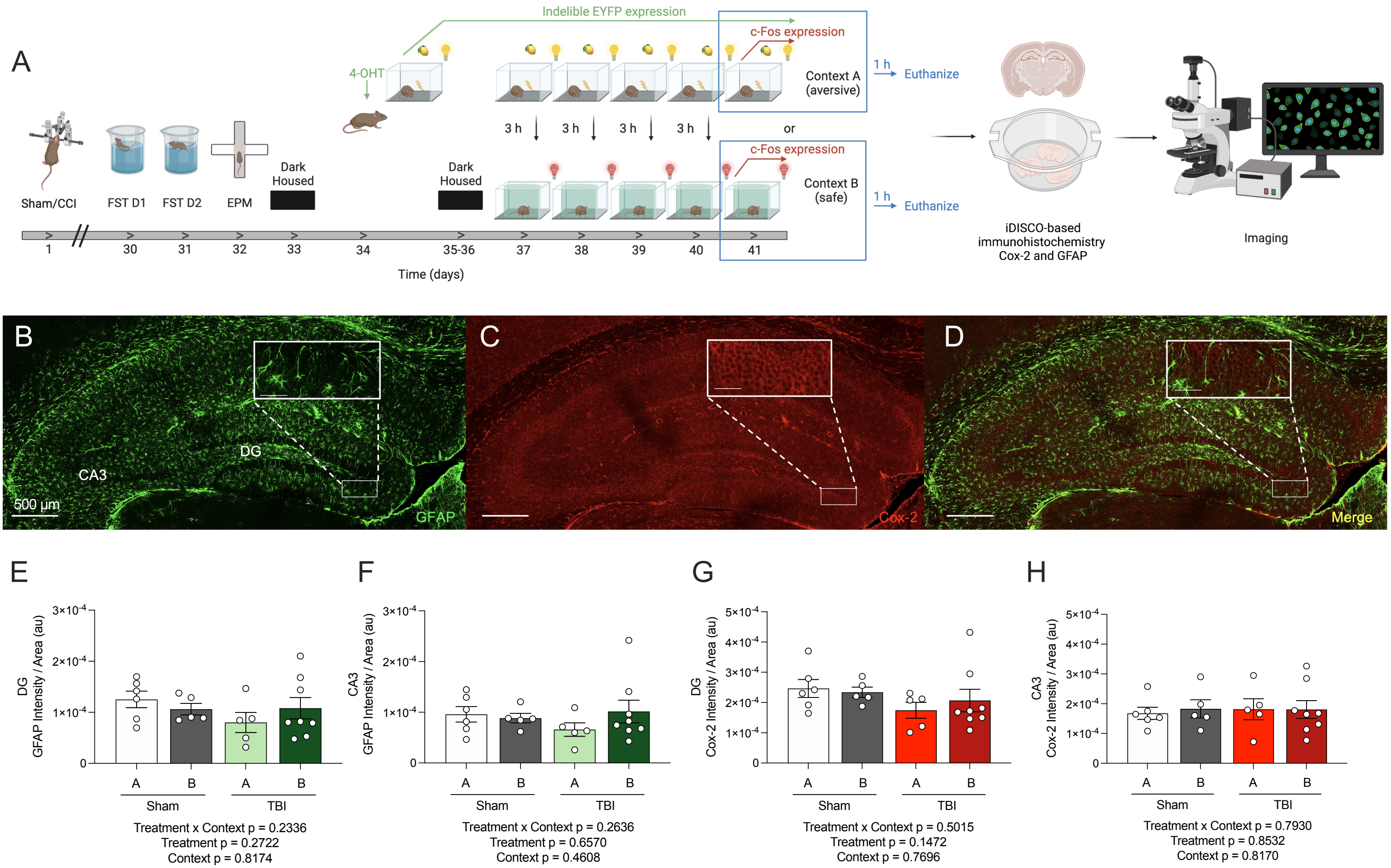

### Figure S4

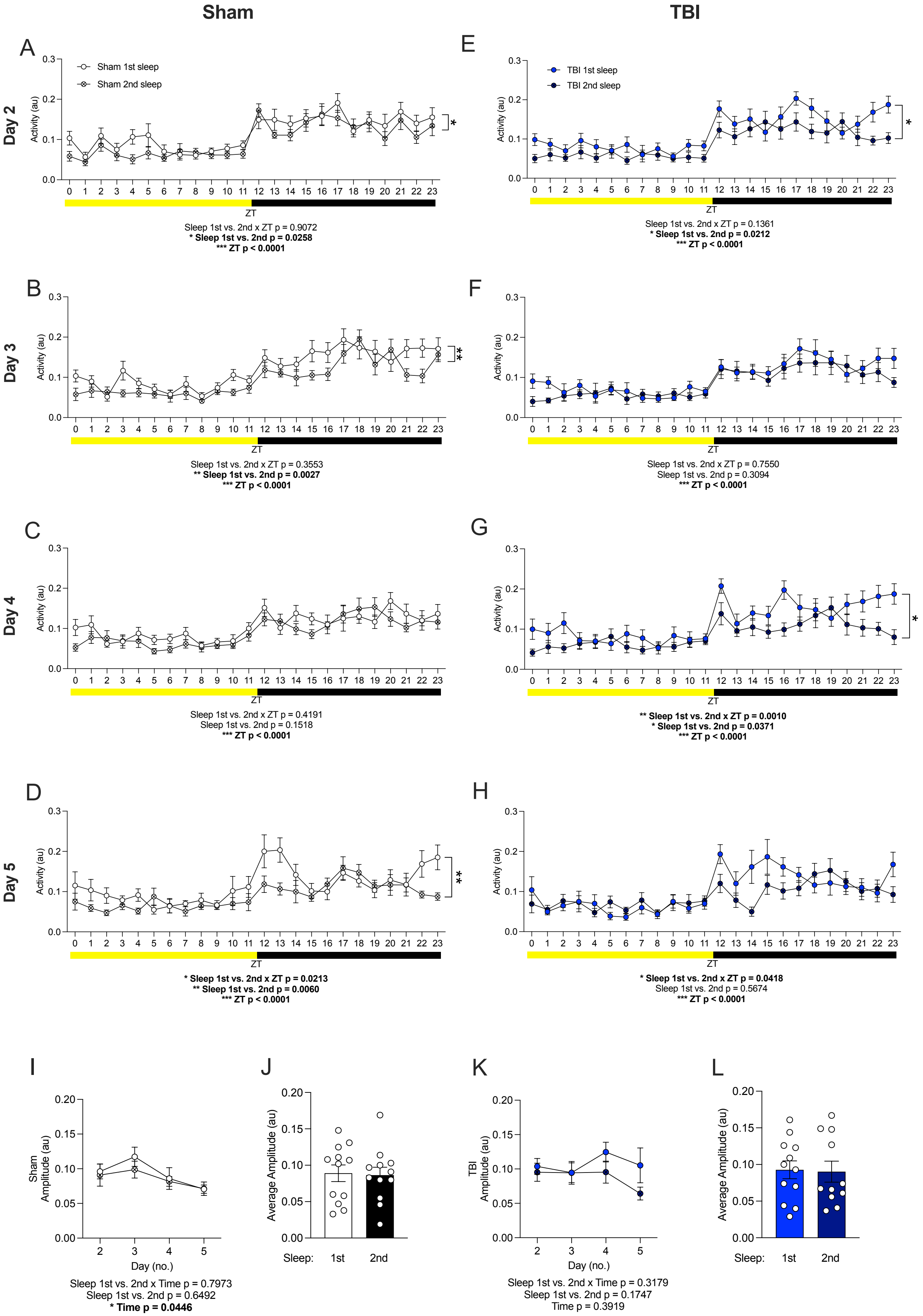

### Figure S5

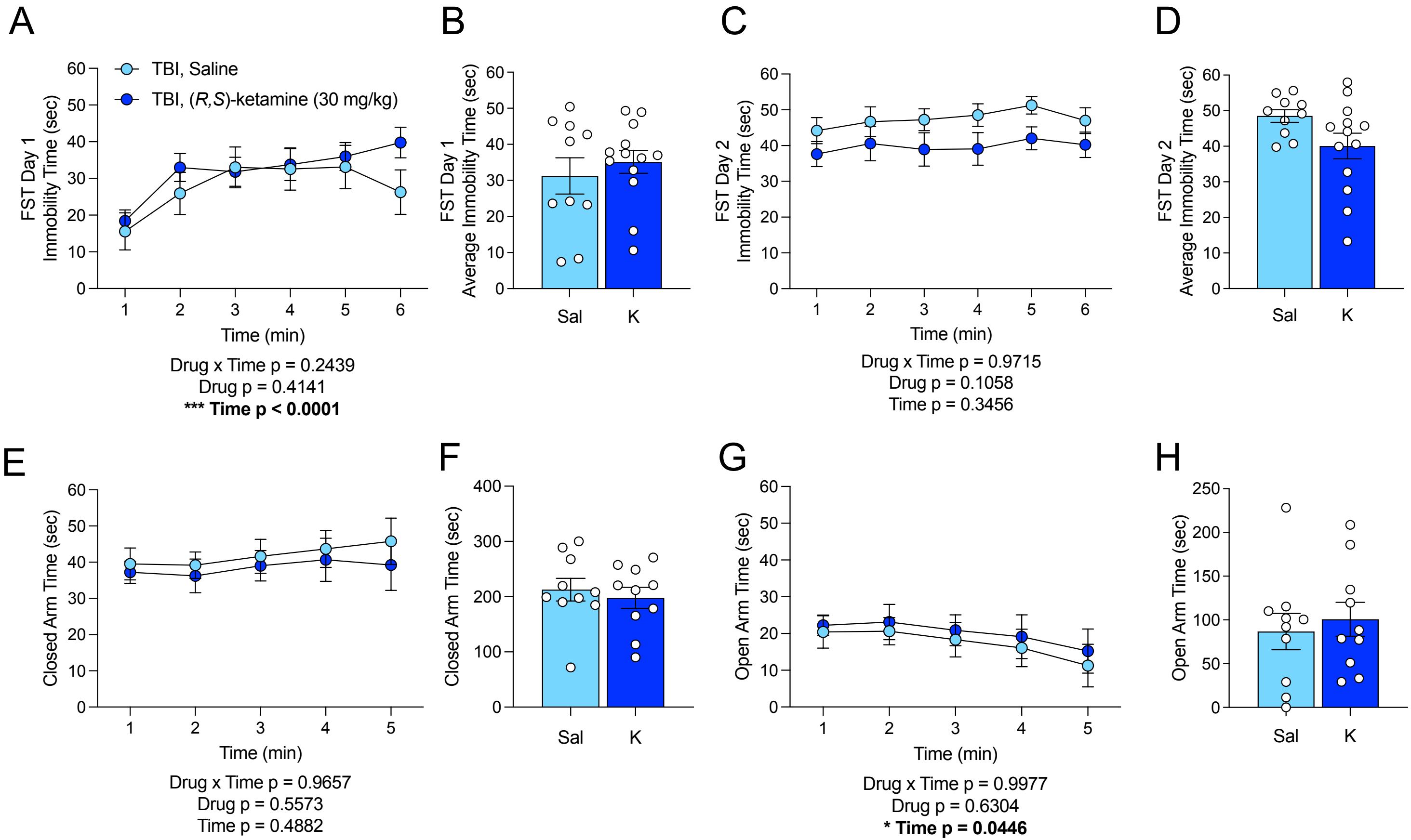
