## Supplementary material for "Traumatic brain injury-induced fear generalization in mice involves hippocampal memory trace dysfunction and is alleviated by (*R,S*)-ketamine": Table S1

**Table 1. Contextual information for contextual fear discrimination contexts.**

| Category | Context A (Aversive Context) | Context B (Neutral Context) |
|---|---|---|
| Overall Context | CFC Chamber | CFC Chamber |
| Shock Administered | One foot shock | None |
| Cleaning Solution | 95% EtOH | Virkon |
| Container Doors | Closed | Open |
| Grids | Exposed | Exposed |
| House Fan | ON | ON |
| House Lights | ON | ON |
| Modified Walls | None | Rounded, colored walls |
| Red Lamps | OFF | ON |
| Room Lights | ON | OFF |
| Scent | Lemon | Anise |
| Transfer Cage | Square Cage | White Bucket |
