## Supplementary material for "Traumatic brain injury-induced fear generalization in mice involves hippocampal memory trace dysfunction and is alleviated by (*R,S*)-ketamine": Table S2

| Cohort | Behavioral Paradigm | Measurement | Statistical Test | Group | Comparison | F or $R^2$ | ° of freedom | p | * | Fig. |
|---|---|---|---|---|---|---|---|---|---|---|
| | | Freezing (%) | 3way RMANOVA | Sham vs. TBI (Treatment) | Treatment x Context x Time | 0.6509 | 3, 112 | 0.5840 | ns | 1D-E |
| | | | | | Treatment x Context | 0.5414 | 3, 112 | 0.4634 | ns | |
| | | | | | Treatment x Time | 0.2414 | 3, 112 | 0.8673 | ns | |
| | | | | | Context x Time | 16.57 | 3, 112 | <0.0001 | **** | |
| | | | | | Treatment | 0.0893 | 1, 112 | 0.7656 | ns | |
| | | | | | Context | 17.79 | 1, 112 | <0.0001 | **** | |
| | | | | | Time | 12.33 | 3, 112 | <0.0001 | **** | |
| | | | 2way RMANOVA | Sham | Context x Time | 6.818 | 4, 121 | 0.0007 | *** | |
| | | | | | Context | 16.28 | 1, 121 | 0.0001 | *** | |
| | | | | | Time | 8.948 | 4, 121 | <0.0001 | **** | |
| | | | | TBI | Context x Time | 2.48 | 4, 151 | 0.0464 | * | |
| | | | | | Context | 3.15 | 1, 151 | 0.0779 | ns | |
| | | | | | Time | 7.623 | 4, 151 | <0.0001 | **** | |
| | | | Sidak's | Sham Day 2 | Context A vs. B | - | - | 0.2565 | ns | |
| | | | | Sham Day 3 | Context A vs. B | - | - | 0.7566 | ns | |
| | | | | Sham Day 4 | Context A vs. B | - | - | 0.0691 | ns | |
| | | | | Sham Day 5 | Context A vs. B | - | - | 0.0046 | ** | |
| | | | | Sham Day 6 | Context A vs. B | - | - | 0.0052 | ** | |
| | | Day 6 Average Freezing (%) | 2way ANOVA | Sham vs. TBI (Treatment) | Treatment x Context | 3.672 | 1, 34 | 0.1076 | ns | 1F |
| | | | | | Treatment | 2.834 | 1, 34 | 0.3663 | ns | |
| | | | | | Context | 5.549 | 1, 34 | 0.0156 | * | |
| | | | Sidak's | Sham | Context A vs. B | - | - | 0.0176 | * | |
| | | | | TBI | Context A vs. B | - | - | 0.7533 | ns | |
| | | EYFP dentate gyrus (DG) Cells per section (No.) | 2way ANOVA | Sham vs. TBI (Treatment) | Treatment x Context | 1.197 | 1, 25 | 0.2843 | ns | 1J |
| | | | | | Treatment | 0.0026 | 1, 25 | 0.9599 | ns | |
| | | | | | Context | 0.4374 | 1, 25 | 0.5144 | ns | |
| | | c-fos DG cells per section (No.) | 2way ANOVA | Sham vs. TBI (Treatment) | Treatment x Context | 2.742 | 1, 25 | 0.1103 | ns | 1K |
| | | | | | Treatment | 0.001 | 1, 25 | 0.9750 | ns | |
| | | | | | Context | 0.0239 | 1, 25 | 0.0856 | ns | |
| | | co-labeled / EYFP DG | 2way ANOVA | Sham vs. TBI (Treatment) | Treatment x Context | 0.2965 | 1, 25 | 0.5909 | ns | 1L |
| | | | | | Treatment | 0.694 | 1, 25 | 0.4127 | ns | |
| | | | | | Context | 11.77 | 1, 25 | 0.0021 | ** | |

| | | | | | | | | | | |
|---|---|---|---|---|---|---|---|---|---|---|
| Memory tagging during Pattern Separation | Pattern Separation (PS) | cells (%) | Sidak's | Sham | Context A vs. B | - | - | 0.0254 | * | |
| | | | | TBI | Context A vs. B | - | - | 0.0812 | ns | |
| | | EYFP CA3 Cells per section (No.) | 2way ANOVA | Sham vs. TBI (Treatment) | Treatment x Context | 0.2261 | 1, 20 | 0.6396 | ns | 2D |
| | | | | | Treatment | 4.455 | 1, 20 | 0.0476 | * | |
| | | | | | Context | 0.6851 | 1, 20 | 0.4176 | ns | |
| | | c-fos CA3 cells per section (No.) | 2way ANOVA | Sham vs. TBI (Treatment) | Treatment x Context | 0.8319 | 1, 20 | 0.3726 | ns | 2E |
| | | | | | Treatment | 7.05 | 1, 20 | 0.0152 | * | |
| | | | | | Context | 2.091 | 1, 20 | 0.1636 | ns | |
| | | co-labeled / EYFP CA3 cells (%) | 2way ANOVA | Sham vs. TBI (Treatment) | Treatment x Context | 0.043 | 1, 20 | 0.8873 | ns | 2F |
| | | | | | Treatment | 4.579 | 1, 20 | 0.0449 | * | |
| | | | | | Context | 0.1417 | 1, 20 | 0.7105 | ns | |
| | | EYFP basal amygdala (BA) Cells per section (No.) | 2way ANOVA | Sham vs. TBI (Treatment) | Treatment x Context | 4.721 | 1, 23 | 0.0404 | * | 3D |
| | | | | | Treatment | 0.0009 | 1, 23 | 0.9770 | ns | |
| | | | | | Context | 0.2703 | 1, 23 | 0.6081 | ns | |
| | | | Sidak's | Sham | Context A vs. B | - | - | 0.1795 | ns | |
| | | | | TBI | Context A vs. B | - | - | 0.3714 | ns | |
| | | c-fos BA cells per section (No.) | 2way ANOVA | Sham vs. TBI (Treatment) | Treatment x Context | 2.224 | 1, 23 | 0.1495 | ns | 3E |
| | | | | | Treatment | 0.6011 | 1, 23 | 0.4461 | ns | |
| | | | | | Context | 2.837 | 1, 23 | 0.1056 | ns | |
| | | co-labeled / EYFP BA cells (%) | 2way ANOVA | Sham vs. TBI (Treatment) | Treatment x Context | 8.83 | 1, 23 | 0.0068 | ** | 3F |
| | | | | | Treatment | 6.057 | 1, 23 | 0.0218 | * | |
| | | | | | Context | 17.01 | 1, 23 | 0.0004 | *** | |
| | | | Sidak's | Sham | Context A vs. B | - | - | 0.0003 | *** | |
| | | | | TBI | Context A vs. B | - | - | 0.6091 | ns | |
| | | EYFP central amygdala (CeA) Cells per section (No.) | 2way ANOVA | Sham vs. TBI (Treatment) | Treatment x Context | 0.3736 | 1, 23 | 0.5470 | ns | 3G |
| | | | | | Treatment | 0.2385 | 1, 23 | 0.6299 | ns | |
| | | | | | Context | 0.0466 | 1, 23 | 0.8310 | ns | |
| | | c-fos CeA cells per section (No.) | 2way ANOVA | Sham vs. TBI (Treatment) | Treatment x Context | 0.0029 | 1, 23 | 0.9574 | ns | 3H |
| | | | | | Treatment | 0.4976 | 1, 23 | 0.4876 | ns | |
| | | | | | Context | 5.251 | 1, 23 | 0.0314 | * | |
| | | | Sidak's | Sham | Context A vs. B | - | - | 0.2637 | ns | |
| | | | | TBI | Context A vs. B | - | - | 0.1759 | ns | |
| | | | 2way | Sham vs. TBI | Treatment x Context | 4.592 | 1, 23 | 0.0429 | * | |

| | | | | | | | | | | |
|---|---|---|---|---|---|---|---|---|---|---|
| | | co-labeled / EYFP CeA cells (%) | 2way ANOVA | Sham vs. TBI (Treatment) | Treatment | 0.544 | 1, 23 | 0.4682 | ns | 3I |
| | | | | | Context | 4.901 | 1, 23 | 0.0370 | * | |
| | | | Sidak's | Sham | Context A vs. B | - | - | 0.0191 | * | |
| | | | | TBI | Context A vs. B | - | - | 0.9981 | ns | |
| | Forced Swim Test (FST) Day 1 | Immobility Time (sec) | 2way RMANOVA | Sham vs. TBI (Treatment) | Treatment x Time | 0.1158 | 5, 140 | 0.9861 | ns | S2A |
| | | | | | Treatment | 16.06 | 1, 28 | 0.7976 | ns | |
| | | | | | Time | 0.0671 | 5, 140 | <0.0001 | *** | |
| | | Average Immobility Time | *t*- test | | Treatment | - | - | 0.7976 | ns | S2B |
| | FST Day 2 | Immobility Time (sec) | 2way RMANOVA | Sham vs. TBI (Treatment) | Treatment x Time | 2.211 | 5, 140 | 0.0565 | ns | S2C |
| | | | | | Treatment | 4.098 | 1, 28 | 0.0526 | ns | |
| | | | | | Time | 10.18 | 5, 140 | <0.0001 | *** | |
| | | Average Immobility Time (min 3-6) | *t*- test | | Treatment | - | - | 0.0498 | * | S2D |
| | Elevated Plus Maze (EPM) | Time in closed arms (sec) | 2way ANOVA | Sham vs. TBI (Treatment) | Treatment x Time | 1.513 | 4, 112 | 0.2032 | ns | S2E |
| | | | | | Treatment | 0.3534 | 1, 28 | 0.5570 | ns | |
| | | | | | Time | 3.262 | 4, 112 | 0.0143 | * | |
| | | | *t*- test | | Treatment | - | - | 0.4306 | ns | S2F |
| | | Time in open arms (sec) | 2way ANOVA | Sham vs. TBI (Treatment) | Treatment x Time | 1.746 | 4, 112 | 0.1448 | ns | S2G |
| | | | | | Treatment | 0.4061 | 1, 28 | 0.5292 | ns | |
| | | | | | Time | 3.242 | 4, 112 | 0.0147 | * | |
| | | | *t*- test | | Treatment | - | - | 0.5416 | ns | S2H |
| Inflammation: GFAP and Cox-2 in DG and CA3 | GFAP | Intensity in DG | 2way ANOVA | Sham vs. TBI (Treatment) | Treatment x Context | 1.509 | 1, 20 | 0.2336 | ns | S3E |
| | | | | | Treatment | 1.275 | 1, 20 | 0.2722 | ns | |
| | | | | | Context | 0.0547 | 1, 20 | 0.8174 | ns | |
| | | Intensity in CA3 | 2way ANOVA | Sham vs. TBI (Treatment) | Treatment x Context | 1.323 | 1, 20 | 0.2636 | ns | S3F |
| | | | | | Treatment | 0.2032 | 1, 20 | 0.6570 | ns | |
| | | | | | Context | 0.5654 | 1, 20 | 0.4608 | ns | |
| | Cox-2 | Intensity in DG | 2way ANOVA | Sham vs. TBI (Treatment) | Treatment x Context | 0.4685 | 1, 20 | 0.5015 | ns | S3G |
| | | | | | Treatment | 2.274 | 1, 20 | 0.1472 | ns | |
| | | | | | Context | 0.0881 | 1, 20 | 0.7696 | ns | |

| | | | | | | | | | | |
|---|---|---|---|---|---|---|---|---|---|---|
| | Cox-2 | Intensity in CA3 | 2way ANOVA | Sham vs. TBI (Treatment) | Treatment x Context | 0.0707 | 1, 20 | 0.7930 | ns | S3H |
| | | | | | Treatment | 0.0352 | 1, 20 | 0.8532 | ns | |
| | | | | | Context | 0.055 | 1, 20 | 0.8170 | ns | |
| | Contextual Fear Conditioning (CFC) Day 2 | Average Freezing (%) | *t*- test | Sham vs. TBI (Treatment) | Treatment | - | - | 0.0366 | * | 4B |
| | Sleep % | Sleep (%) Total | 2way RMANOVA | 1st vs. 2nd sleep | Sleep 1st vs. 2nd x Treatment | 0.1337 | 1, 42 | 0.1337 | ns | 4C |
| | | | | | 1st vs. 2nd sleep | 0.0891 | 1, 42 | 0.0891 | ns | |
| | | | | | Treatment | 0.2134 | 1, 42 | 0.2134 | ns | |
| | | Sleep (%) Day | 2way RMANOVA | 2nd vs. 2nd sleep | Sleep 1st vs. 2nd x Treatment | 2.16 | 1, 42 | 0.1491 | ns | 4D |
| | | | | | 1st vs. 2nd sleep | 1.012 | 1, 42 | 0.3201 | ns | |
| | | | | | Treatment | 4.834 | 1, 42 | 0.0335 | * | |
| | | | Sidak's | 1st sleep | Treatment | - | - | 0.0258 | * | |
| | | | | 2nd sleep | Treatment | - | - | 0.8471 | ns | |
| | | Sleep (%) Night | 2way RMANOVA | 1st vs. 2nd sleep | Sleep 1st vs. 2nd x Treatment | 0.7694 | 1, 42 | 0.3854 | ns | 4E |
| | | | | | Sham 1st vs. 2nd sleep | | 1, 42 | 0.3748 | ns | |
| | | | | | Treatment | | 1, 42 | 0.5763 | ns | |
| | | Sleep (%) Total 2nd sleep and CFC Day 2 Freezing (%) | Pearson's Correlation | Sham | - | 0.6609 | - | 0.0013 | ** | 4F |
| | | | | TBI | - | 0.0171 | - | 0.7019 | ns | |
| | Sleep Bout | Average Sleep Bout Length | 2way RMANOVA | 1st vs. 2nd sleep | Sleep 1st vs. 2nd x Treatment | 0.1268 | 1, 42 | 0.2558 | ns | 4G |
| | | | | | 1st vs. 2nd sleep x ZT | 6.417 | 1, 42 | 0.0151 | * | |
| | | | | | Treatment | 1.096 | 1, 42 | 0.3011 | ns | |
| | Amplitude | Average amplitude (au) | 2way RMANOVA | 1st vs. 2nd sleep | Sleep 1st vs. 2nd x Treatment | 1.891 x 10^-5 | 1, 43 | 0.9966 | ns | 4H |
| | | | | | 1st vs. 2nd sleep | 0.0431 | 1, 43 | 0.8366 | ns | |
| | | | | | Treatment | 0.0972 | 1, 43 | 0.7567 | ns | |
| | | | 3way RMANOVA | Sham vs. TBI (Treatment) | Treatment x 1st vs. 2nd sleep x ZT | 0.6482 | 2, 42 | 0.5281 | ns | 4I-J |
| | | | | | 1st vs. 2nd sleep x ZT | 0.4558 | 2, 42 | 0.637 | ns | |
| | | | | | Treatment x ZT | 0.6957 | 2, 42 | 0.5044 | ns | |
| | | | | | Treatment x 1st vs. 2nd sleep | 0.5745 | 1, 21 | 0.4569 | * | |

| | | | | | | | | | | |
|---|---|---|---|---|---|---|---|---|---|---|
| | | | | | Treatment | 0.0009 | 1, 21 | 0.9763 | ns | |
| | | | | | ZT | 73.73 | 2, 42 | <0.0001 | **** | |
| | | | | | 1st vs. 2nd sleep | | 1, 21 | 0.0443 | * | |
| | | | 2way RMANOVA | Sham 1st vs. 2nd sleep | Sleep 1st vs. 2nd x ZT | 1.067 | 23, 506 | 0.3785 | ns | |
| | | | | | Sleep 1st vs. 2nd | 11.63 | 1, 22 | 0.0025 | ** | |
| | | | | | ZT | 1.067 | 7.733, 170.1 | <0.0001 | **** | |
| | | | | ZT0 | Sham 1st vs. 2nd sleep | - | - | 0.0283 | * | |
| | | | | ZT1 | Sham 1st vs. 2nd sleep | - | - | 0.0483 | * | |
| | | | | ZT2 | Sham 1st vs. 2nd sleep | - | - | 0.2318 | ns | |
| | | | | ZT3 | Sham 1st vs. 2nd sleep | - | - | 0.0407 | * | |
| | | | | ZT4 | Sham 1st vs. 2nd sleep | - | - | 0.0034 | ** | |
| | | | | ZT5 | Sham 1st vs. 2nd sleep | - | - | 0.227 | ns | |
| | | | | ZT6 | Sham 1st vs. 2nd sleep | - | - | 0.172 | ns | |
| | | | | ZT7 | Sham 1st vs. 2nd sleep | - | - | 0.1796 | ns | |
| | | | | ZT8 | Sham 1st vs. 2nd sleep | - | - | 0.184 | ns | |
| | | | | ZT9 | Sham 1st vs. 2nd sleep | - | - | 0.1385 | ns | |
| | | | | ZT10 | Sham 1st vs. 2nd sleep | - | - | 0.1369 | ns | 4I |
| | | | | ZT11 | Sham 1st vs. 2nd sleep | - | - | 0.112 | ns | |
| | | | Sidak's | ZT12 | Sham 1st vs. 2nd sleep | - | - | 0.1029 | ns | |
| | | | | ZT13 | Sham 1st vs. 2nd sleep | - | - | 0.0875 | ns | |
| | | | | ZT14 | Sham 1st vs. 2nd sleep | - | - | 0.1001 | ns | |
| | | | | ZT15 | Sham 1st vs. 2nd sleep | - | - | 0.0493 | * | |
| | | | | ZT16 | Sham 1st vs. 2nd sleep | - | - | 0.1811 | ns | |
| | | | | ZT17 | Sham 1st vs. 2nd sleep | - | - | 0.1391 | ns | |
| | | | | ZT18 | Sham 1st vs. 2nd sleep | - | - | 0.2331 | ns | |
| | Average activity over time | Average activity (au) | | ZT19 | Sham 1st vs. 2nd sleep | - | - | 0.6824 | ns | |
| | | | | ZT20 | Sham 1st vs. 2nd sleep | - | - | 0.3951 | ns | |
| | | | | ZT21 | Sham 1st vs. 2nd sleep | - | - | 0.0362 | * | |
| | | | | ZT22 | Sham 1st vs. 2nd sleep | - | - | 0.0272 | * | |
| | | | | ZT23 | Sham 1st vs. 2nd sleep | - | - | 0.1024 | ns | |
| | | | 2way RMANOVA | TBI 1st vs. 2nd sleep | Sleep 1st vs. 2nd x ZT | 1.717 | 22, 483 | 0.0210 | * | |
| | | | | | Sleep 1st vs. 2nd | 1.859 | 1, 21 | 0.1872 | ns | |
| | | | | | ZT | 24.53 | 6.764, 142.1 | <0.0001 | *** | |
| | | | | ZT0 | TBI 1st vs. 2nd sleep | - | - | 0.0172 | * | |

| | | | | | | | | | | |
|---|---|---|---|---|---|---|---|---|---|---|
| Sleep | | | Sidak's | ZT1 | TBI 1st vs. 2nd sleep | - | - | 0.045 | * | 4J |
| | | | | ZT2 | TBI 1st vs. 2nd sleep | - | - | 0.1645 | ns | |
| | | | | ZT3 | TBI 1st vs. 2nd sleep | - | - | 0.3608 | ns | |
| | | | | ZT4 | TBI 1st vs. 2nd sleep | - | - | 0.9362 | ns | |
| | | | | ZT5 | TBI 1st vs. 2nd sleep | - | - | 0.1639 | ns | |
| | | | | ZT6 | TBI 1st vs. 2nd sleep | - | - | 0.3004 | ns | |
| | | | | ZT7 | TBI 1st vs. 2nd sleep | - | - | 0.7565 | ns | |
| | | | | ZT8 | TBI 1st vs. 2nd sleep | - | - | 0.1974 | ns | |
| | | | | ZT9 | TBI 1st vs. 2nd sleep | - | - | 0.4072 | ns | |
| | | | | ZT10 | TBI 1st vs. 2nd sleep | - | - | 0.2492 | ns | |
| | | | | ZT11 | TBI 1st vs. 2nd sleep | - | - | 0.5716 | ns | |
| | | | | ZT12 | TBI 1st vs. 2nd sleep | - | - | 0.2242 | ns | |
| | | | | ZT13 | TBI 1st vs. 2nd sleep | - | - | 0.7026 | ns | |
| | | | | ZT14 | TBI 1st vs. 2nd sleep | - | - | 0.503 | ns | |
| | | | | ZT15 | TBI 1st vs. 2nd sleep | - | - | 0.6587 | ns | |
| | | | | ZT16 | TBI 1st vs. 2nd sleep | - | - | 0.3924 | ns | |
| | | | | ZT17 | TBI 1st vs. 2nd sleep | - | - | 0.1096 | ns | |
| | | | | ZT18 | TBI 1st vs. 2nd sleep | - | - | 0.973 | ns | |
| | | | | ZT19 | TBI 1st vs. 2nd sleep | - | - | 0.3135 | ns | |
| | | | | ZT20 | TBI 1st vs. 2nd sleep | - | - | 0.6917 | ns | |
| | | | | ZT21 | TBI 1st vs. 2nd sleep | - | - | 0.3911 | ns | |
| | | | | ZT22 | TBI 1st vs. 2nd sleep | - | - | 0.0942 | ns | |
| | | | | ZT23 | TBI 1st vs. 2nd sleep | - | - | 0.0013 | ** | |
| | Activity during ZT11 | Activity (au) | 2way ANOVA | Sham 1st vs. 2nd sleep | Sleep 1st vs. 2nd x Treatment | 0.4161 | 1, 42 | 0.5224 | ns | 4K |
| | | | | | Sleep 1st vs. 2nd | 2.529 | 1, 42 | 0.1193 | ns | |
| | | | | | Treatment | 0.375 | 1, 42 | 0.5436 | ns | |
| | Activity during ZT23 | Activity (au) | 2way ANOVA | Sham 1st vs. 2nd sleep | Sleep 1st vs. 2nd x Treatment | 1.141 | 1, 43 | 0.2914 | ns | 4L |
| | | | | | Sleep 1st vs. 2nd | 13.44 | 1, 43 | 0.0007 | *** | |
| | | | | | Treatment | 0.3925 | 1, 43 | 0.5343 | ns | |
| | | | Sidak's | Sham | 1st vs. 2nd sleep | - | - | 0.1352 | ns | |
| | | | | TBI | 1st vs. 2nd sleep | - | - | 0.0038 | ** | |
| | | Day 2 | 2way RMANOVA | 1st vs. 2nd sleep | 1st vs. 2nd sleep x ZT | 0.6324 | 23, 506 | 0.9072 | ns | S4A |
| | | | | | 1st vs. 2nd sleep | 5.712 | 1, 22 | 0.0258 | * | |
| | | | | | ZT | 9.819 | 8.803, 193.7 | <0.0001 | **** | |

| | | | | | | | | | | |
|---|---|---|---|---|---|---|---|---|---|---|
| | Activity over time - Sham | Day 3 | 2way RMANOVA | 1st vs. 2nd sleep | 1st vs. 2nd sleep x ZT | 1.087 | 23, 506 | 0.3553 | ns | S4B |
| | | | | | 1st vs. 2nd sleep | 11.39 | 1, 22 | 0.0027 | ** | |
| | | | | | ZT | 10.96 | 8.104, 178.3 | <0.0001 | *** | |
| | | Day 4 | Mixed-effects 2way RMANOVA | 1st vs. 2nd sleep | 1st vs. 2nd sleep x ZT | 1.035 | 23, 470 | 0.4191 | ns | S4C |
| | | | | | 1st vs. 2nd sleep | 2.205 | 1, 22 | 0.1518 | ns | |
| | | | | | ZT | 7.137 | 9.159, 187.2 | <0.0001 | *** | |
| | | Day 5 | Mixed-effects 2way RMANOVA | 1st vs. 2nd sleep | 1st vs. 2nd sleep x ZT | 1.727 | 23, 344 | 0.0213 | * | S4D |
| | | | | | 1st vs. 2nd sleep | 9.709 | 1, 18 | 0.006 | ** | |
| | | | | | ZT | 4.945 | 6.769, 84.17 | <0.0001 | *** | |
| | Activity over time - TBI | Day 2 | 2way RMANOVA | 1st vs. 2nd sleep | 1st vs. 2nd sleep x ZT | 1.338 | 23, 506 | 0.1361 | ns | S4E |
| | | | | | 1st vs. 2nd sleep | 6.164 | 1, 22 | 0.0212 | * | |
| | | | | | ZT | 10.47 | 9.577, 210.7 | <0.0001 | *** | |
| | | Day 3 | 2way RMANOVA | 1st vs. 2nd sleep | 1st vs. 2nd sleep x ZT | 0.7824 | 23, 506 | 0.755 | ns | S4F |
| | | | | | 1st vs. 2nd sleep | 1.083 | 1, 22 | 0.3094 | ns | |
| | | | | | ZT | 10.9 | 8.078, 177.7 | <0.0001 | *** | |
| | | Day 4 | Mixed-effects 2way RMANOVA | 1st vs. 2nd sleep | 1st vs. 2nd sleep x ZT | 2.225 | 23, 470 | 0.001 | ** | S4G |
| | | | | | 1st vs. 2nd sleep | 4.926 | 1, 22 | 0.0371 | * | |
| | | | | | ZT | 9.256 | 7.717, 157.7 | <0.0001 | **** | |
| | | Day 5 | Mixed-effects 2way RMANOVA | 1st vs. 2nd sleep | 1st vs. 2nd sleep x ZT | 1.597 | 23, 344 | 0.0418 | * | S4H |
| | | | | | 1st vs. 2nd sleep | 0.3394 | 1, 18 | 0.5674 | ns | |
| | | | | | ZT | 7.176 | 6.344, 94.89 | <0.0001 | **** | |
| | Amplitude | Amplitude (au) over time | Mixed-effects 2way RMANOVA | 1st vs. 2nd sleep | 1st vs. 2nd sleep x Time | 0.3389 | 3, 50 | 0.7973 | ns | S4I |
| | | | | | 1st vs. 2nd sleep | 0.2127 | 1, 22 | 0.6492 | ns | |
| | | | | | Time | 2.995 | 2.739, 45.65 | 0.0446 | * | |
| | | Average amplitude (au) | *t*-test | Sham | 1st vs. 2nd sleep | - | - | 0.8306 | ns | S4J |

| | | | | | | | | | | |
|---|---|---|---|---|---|---|---|---|---|---|
| | Amplitude | Amplitude (au) over time | Mixed-effects 2way RMANOVA | 1st vs. 2nd sleep | 1st vs. 2nd sleep x Time | 1.206 | 3, 47 | 0.3179 | ns | S4K |
| | | | | | 1st vs. 2nd sleep | 1.973 | 1, 21 | 0.1747 | ns | |
| | | | | | Time | 0.9877 | 2.294, 35.94 | 0.3939 | ns | |
| | | Average amplitude (au) | *t*-test | TBI | 1st vs. 2nd sleep | - | - | 0.7034 | ns | S4L |
| | PS | Freezing (%) | Mixed-effects 3way RMANOVA | Saline vs. (*R,S*)-ketamine (Drug) | Drug x Context x Time | 0.7717 | 8, 154 | 0.6282 | ns | 5B-C |
| | | | | | Drug x Context | 0.116 | 1, 154 | 0.7338 | ns | |
| | | | | | Drug x Time | 0.2939 | 8, 161 | 0.9672 | ns | |
| | | | | | Context x Time | 7.478 | 8, 154 | <0.0001 | *** | |
| | | | | | Drug | 15.99 | 1, 161 | <0.0001 | *** | |
| | | | | | Context | 27.2 | 1, 154 | <0.0001 | *** | |
| | | | | | Time | 1.413 | 8, 161 | 0.1948 | ns | |
| | | | Mixed-effects 2way RMANOVA | Saline | Context x Time | 1.413 | 8,99 | 0.2003 | ns | |
| | | | | | Context | 2.652 | 1, 18 | 0.1208 | ns | |
| | | | | | Time | 1.187 | 8, 99 | 0.3146 | ns | |
| | | | | (*R,S*)-ketamine | Context x Time | 8.035 | 8, 174 | <0.0001 | **** | |
| | | | | | Context | 3.063 | 1, 24 | 0.0929 | ns | |
| | | | | | Time | 2.577 | 5.5, 120.1 | 0.0254 | * | |
| | | | Sidak's | Sal Day 2 | Context A vs. B | - | - | >0.9999 | ns | |
| | | | | Sal Day 3 | Context A vs. B | - | - | 0.9999 | ns | |
| | | | | Sal Day 4 | Context A vs. B | - | - | >0.9999 | ns | |
| | | | | Sal Day 5 | Context A vs. B | - | - | 0.9677 | ns | |
| | | | | Sal Day 6 | Context A vs. B | - | - | 0.9816 | ns | |
| | | | | Sal Day 7 | Context A vs. B | - | - | 0.8585 | ns | |
| | | | | Sal Day 8 | Context A vs. B | - | - | 0.7559 | ns | |
| | | | | Sal Day 9 | Context A vs. B | - | - | 0.4835 | ns | |
| | | | | Sal Day 10 | Context A vs. B | - | - | 0.2584 | ns | |
| | | | | K Day 2 | Context A vs. B | - | - | 0.1191 | ns | |
| | | | | K Day 3 | Context A vs. B | - | - | 0.3048 | ns | |
| | | | | K Day 4 | Context A vs. B | - | - | >0.9999 | ns | |
| | | | | K Day 5 | Context A vs. B | - | - | 0.8425 | ns | |
| | | | | K Day 6 | Context A vs. B | - | - | 0.9931 | ns | |
| | | | | K Day 7 | Context A vs. B | - | - | 0.7677 | ns | |

| | | | | | | | | | | |
|---|---|---|---|---|---|---|---|---|---|---|
| (*R,S*)-ketamine post-impact | | | | K Day 8 | Context A vs. B | - | - | 0.8345 | ns | |
| | | | | K Day 9 | Context A vs. B | - | - | 0.0306 | * | |
| | | | | K Day 10 | Context A vs. B | - | - | 0.0004 | *** | |
| | | Day 10 Average Freezing (%) | 2way ANOVA | Saline vs. (*R,S*)-ketamine (Drug) | Drug x Context | 2.703 | 1, 42 | 0.1076 | ns | 5D |
| | | | | | Drug | 3.028 | 1, 42 | 0.0892 | ns | |
| | | | | | Context | 18.51 | 1, 42 | <0.0001 | *** | |
| | | | Sidak's | Saline | Context A vs. B | - | - | 0.1614 | ns | |
| | | | | (*R,S*)-ketamine | Context A vs. B | - | - | 0.0001 | *** | |
| | | EYFP dentate gyrus (DG) Cells per section (No.) | 2way ANOVA | Saline vs. (*R,S*)-ketamine (Drug) | Drug x Context | 2.961 | 1, 19 | 0.1015 | ns | 5H |
| | | | | | Drug | 2.254 | 1, 19 | 0.1497 | ns | |
| | | | | | Context | 1.822 | 1, 19 | 0.1930 | ns | |
| | | c-fos DG cells per section (No.) | 2way ANOVA | Saline vs. (*R,S*)-ketamine (Drug) | Drug x Context | 0.0001 | 1, 19 | 0.9905 | ns | 5I |
| | | | | | Drug | 0.0351 | 1, 19 | 0.8535 | ns | |
| | | | | | Context | 7.277 | 1, 19 | 0.0143 | * | |
| | | | Sidak's | Sham | Context A vs. B | - | - | 0.1608 | ns | |
| | | | | TBI | Context A vs. B | - | - | 0.1152 | ns | |
| | | co-labeled / EYFP DG cells (%) | 2way ANOVA | Saline vs. (*R,S*)-ketamine (Drug) | Drug x Context | 2.857 | 1, 19 | 0.1073 | ns | 5J |
| | | | | | Drug | 2.284 | 1, 19 | 0.1472 | ns | |
| | | | | | Context | 7.311 | 1, 19 | 0.0141 | * | |
| | | | Sidak's | Sham | Context A vs. B | - | - | 0.5033 | ns | |
| | | | | TBI | Context A vs. B | - | - | 0.0039 | ** | |
| | Forced Swim Test (FST) Day 1 | Immobility Time (sec) | 2way RMANOVA | Saline vs. (*R,S*)-ketamine (Drug) | Drug x Time | 1.364 | 5, 105 | 0.2439 | ns | S5A |
| | | | | | Drug | 0.6941 | 1, 21 | 0.4141 | ns | |
| | | | | | Time | 8.63 | 5, 105 | <0.0001 | *** | |
| | | Average Immobility Time | *t*-test | | Drug | - | - | 0.4996 | ns | S5B |
| | | Immobility Time (sec) | 2way RMANOVA | Saline vs. (*R,S*)- | Drug x Time | 0.1747 | 5, 105 | 0.9715 | ns | S5C |
| | | | | | Drug | 2.857 | 1, 21 | 0.1058 | ns | |
| | | | | | Time | 1.137 | 5, 105 | 0.3456 | ns | |

| | | | | | | | | | | |
|---|---|---|---|---|---|---|---|---|---|---|
| | FST Day 2 | Average Immobility Time (min 3-6) | *t*- test | (*R,S*)-ketamine (Drug) | Drug | - | - | 0.0695 | ns | S5D |
| | Elevated Plus Maze (EPM) | Time in closed arms (sec) | 2way ANOVA | Saline vs. (*R,S*)-ketamine (Drug) | Drug x Time | 0.1426 | 4, 72 | 0.9657 | ns | S5E |
| | | | | | Drug | 0.3575 | 1, 18 | 0.5573 | ns | |
| | | | | | Time | 0.8667 | 4, 72 | 0.4882 | ns | |
| | | | *t*- test | | Drug | - | - | 0.5997 | ns | S5F |
| | | Time in open arms (sec) | 2way ANOVA | Saline vs. (*R,S*)-ketamine (Drug) | Drug x Time | 0.0346 | 4,72 | 0.9977 | ns | S5G |
| | | | | | Drug | 0.2396 | 1.18 | 0.6304 | ns | |
| | | | | | Time | 2.577 | 4,72 | 0.0446 | * | |
| | | | *t*- test | | Drug | - | - | 0.6277 | ns | S5H |
