## Supplementary material for "Traumatic brain injury-induced fear generalization in mice involves hippocampal memory trace dysfunction and is alleviated by (*R,S*)-ketamine": Table S3

**Supplemental Table 3.**

KEY RESOURCES TABLE

| Resource Type | Specific Reagent or Resources | Source or Reference | Identifiers | Additional Information |
|---|---|---|---|---|
| Add additional rows as needed for each resource type | Include species and sex when applicable. | Include name of manufacturer, company, repository, individual, or research lab. Include PMID or DOI for references; use "this paper" if new. | Include catalog numbers, stock numbers, database IDs or accession numbers, and/or RRIDs. RRIDs are highly encouraged; search for RRIDs at https://scicrunch.org/resources | Include any additional information or notes if necessary |
| Antibody | Rabbit polyclonal IgG anti-c-fos | Santa Cruz Biotechnology Inc., Dallas, TX | Cat.# sc-166940 | concentration 1:1000 |
| Antibody | Chicken polyclonal anti-GFP | Abcam, Cambridge, MA | Cat.# ab13970 | concentration 1:2000 |
| Antibody | Cy2 conjugated Donkey Anti-Chicken IgG | Jackson ImunoResearch, West Grove, PA | Cat.# 703-225-155, RRID: AB_2340370 | concentration 1:250 |
| Antibody | Alexa 647 conjugated Donkey Anti-Rabbit IgG | Life Technologies, Carlsbad, CA | Cat.# A-31573, RRID: AB_2536183 | concentration 1:500 |
| Antibody | Mouse monoclonal IgG anti-Cox-2 | Santa Cruz Biotechnology, Santa Cruz, CA | Cat.# sc-19999 | concentration 1:200 |
| Antibody | Rabbit polyclonal IgG anti-GFAP | Agilent Dako Omnis, Santa Clara, CA | Cat.# GA52461-2 | concentration 1:500 |
| Antibody | Alexa 594 conjugated Donkey Anti-Mouse IgG | Thermo Fisher Scientific, Waltham, MA | Cat.# R37115, RRID: AB_2556543 | concentration 1:500 |
| Antibody | Cy2 conjugated Donkey Anti-Rabbit IgG | Jackson ImmunoResearch, West Grove, PA | Cat.# 711-225-152, RRID: AB_2340612 | concentration 1:250 |
| Mounting medium | Fluoromount G | Electron Microscopy Sciences, Hatfield, PA | Cat.# 17984-25 | |
| Organism / strain | Mouse: ArcCreER$^{T2}$(+) x EYFP | doi: 10.1016/j.neuron.2014.05.018 | Non applicable | |
| Chemical Compound or Drug | (*R*,*S*)-ketamine | Fort Dodge Animal Health, Fort Dodge, IA | Cat.# 0856-4403-01 | 30 mg/kg, i.p. |
| Chemical Compound or Drug | 4-Hydroxytamoxifen | Sigma-Aldrich, St. Louis, MO | Cat.# H6278 | |
| Software; Algorithm | ImageJ v.1.52p | http://fiji.sc/ | | |
| Software; Algorithm | ClockLab | http://actimetrics.com/products/clocklab | | |
